# Emergence and function of cortical offset responses in sound termination detection

**DOI:** 10.1101/2021.04.13.439639

**Authors:** Magdalena Sołyga, Tania Rinaldi Barkat

**Affiliations:** Brain & Sound Lab, Department of Biomedicine, Basel University, 4056 Basel, Switzerland

## Abstract

Offset responses in auditory processing appear after a sound terminates. They arise in neuronal circuits within the peripheral auditory system, but their role in the central auditory system remains unknown. Here we ask what the behavioural relevance of cortical offset responses is and what circuit mechanisms drive them. At the perceptual level, our results reveal that experimentally minimizing auditory cortical offset responses decreases the mouse performance to detect sound termination, assigning a behavioural role to offset responses. By combining *in vivo* electrophysiology in the auditory cortex and thalamus of awake mice, we also demonstrate that cortical offset responses are not only inherited from the periphery but also amplified and generated *de novo*. Finally, we show that offset responses code more than silence, including relevant changes in sound trajectories. Together, our results reveal the importance of cortical offset responses in encoding sound termination and detecting changes within temporally discontinuous sounds crucial for speech and vocalization.

## INTRODUCTION

Offset responsive neurons are present through the whole auditory pathway starting from the cochlear nucleus (CN) (Ding et al., 1999; Suga, 1964; Young and Brownell, 1976) to the superior paraolivary nucleus (SPN) (Dehmel et al., 2002; Kulesza et al., 2003), the inferior colliculus (IC) (Akimov et al., 2017; Kasai et al., 2012), the medial geniculate body (MGB) (Anderson and Linden, 2016; He, 2001, 2002; Yu et al., 2004) and the auditory cortex (ACx) (Qin et al., 2007; Recanzone, 2000; Scholl et al., 2010; Takahashi et al., 2004). Multiple mechanisms were proposed as contributing to producing offset responses (Bondanelli et al., 2019; Kopp-Scheinpflug et al., 2018; Xu et al., 2014). Generally it is thought that signals from the cochlea can generate offset responses in both the CN (Suga, 1964) and the SPN (Kopp-Scheinpflug et al., 2011), but strong offset responses were mainly described in the SPN. This structure is considered to be specialized for offset response generation based on the strong inhibitory signal it receives during the sound and on the precise firing when sound ends. The SPN sends then strong inhibitory inputs to the IC (Kulesza and Berrebi, 2000; Saldana et al., 2009), which in turn might further convey the signal to the MGB. Offset responses in the MGB and in the ACx are generally thought to be driven by excitatory/inhibitory inputs from IC rather than by other neural mechanisms (Kopp-Scheinpflug et al., 2018). *De novo* generation or amplification of offset responses in these areas have not been clearly demonstrated yet (Bondanelli et al., 2019; He, 2003; Kasai et al., 2012; Yu et al., 2004).

The perceptual significance of offset responses has been difficult to assess. They have been postulated to play a role in sound duration selectivity (He, 2002; Qin et al., 2009), in gap detection (Syka et al., 2002; Threlkeld et al., 2008; Weible et al., 2014a; Weible et al., 2014b) or in perceiving communication calls (Eggermont, 2015; Felix et al., 2018; Kopp-Scheinpflug et al., 2018). For example, Qin et al. showed that onset-only neurons in the primary ACx of cats cannot discriminate duration, and suggested that sustained and offset responses underlie discrimination of sound duration(Qin et al., 2009). In another study, Weible et al. demonstrated that the cortical postgap neural activity in mice is related to detecting brief gaps in noise (Weible et al., 2014b), but the relative contributions of sound offset and onset responses are unclear (Kopp-Scheinpflug et al., 2018). No evidence has yet demonstrated whether the increased neuronal activity of sound offset responses accounts for these perceptual skills. Compared to onset responses, offset responses in the central auditory pathway are typically less prevalent (Phillips et al., 2002; Pollak and Bodenhamer, 1981; Solyga and Barkat, 2019). They are generally weaker and slower than onset responses (Qin et al., 2007). At the cortical level, offset responses have been shown to cluster within the anterior auditory field (AAF) - a primary region of the ACx - where they have been observed in twice as many cells as in the primary auditory cortex (A1) (Solyga and Barkat, 2019). Offset responses are also highly influenced by different sounds parameters. For example, the amplitude, duration, frequency, fall time and spectral complexity of the sound have all been reported to influence auditory offset responses (He, 2002; Scholl et al., 2010; Solyga and Barkat, 2019; Takahashi et al., 2004). However, no study has yet addressed these influences in a systematic way. The involvement of the different nuclei of the central auditory system, as well as their cellular and circuit mechanisms, are thus poorly understood.

We combined behavioural experiments with optogenetics, electrophysiological recordings and antidromic stimulation to better understand the role that cortical offset responses play in sound perception and the properties that distinguish them from the subcortical ones. Our results reveal that the AAF is highly specialized for processing information on sound termination and that minimizing its offset responses decreases the mouse performance to detect when a sound ends. By studying the influence of different sound parameters on AAF and MGB offset responses, we demonstrate that cortical offsets are inherited, amplified and sometimes even generated *de novo*. First, we found that the AAF, unlike the MGB, shows a significant increase in offset response amplitudes with sound duration. Then, we report that white noise bursts - unlike pure tones - only evoke offset responses in the AAF but not in the MGB. Finally, we show that offset responses are present in the AAF whenever a frequency component is removed from multi frequency stimuli and therefore may have a further role than solely coding silence.

Taken together, our findings suggest a particular involvement of AAF offset responses in sound termination processing and point to the importance of this cortical subfield for advanced processing such as tracking sound duration or detecting changes in frequency and level within temporally discontinuous sounds.

## RESULTS

### Cortical offset responses improve sound termination detection

The perceptual significance of cortical offset responses has been difficult to assess. Indeed, confounds about perceiving a sound and its termination are intricately linked, and changing the neuronal activity of sound offset responses without changing any other parameters of the sound response has not been tested. To assess the behavioural relevance of auditory offset responses, we developed a sound termination detection task in which mice expressing Channelrhodopsin-2 (ChR2) in parvalbumin-positive (PV+) cells learned to detect the end of 9 kHz frequency pure tones (Figure 1a). Animals were placed in a cardboard tube with a speaker 10 cm away from their left ear. A piezo sensor attached to a licking spout was used to measure sound termination detection during the task. To accelerate the learning process, mild air puffs were given when mice licked to sound onset. The duration of the tones was varied randomly (1, 1.5 or 2 s) to avoid a putative expected behavioural response at a fixed delay after sound onset. Mice were initially trained with a tone played at 80 dB SPL. A reaction time window was set to 3 s during initial training sessions and was progressively shorten to 1s in the final training sessions. An increase in correct offset detection (hit rate), and a decrease in reaction time over the training sessions, reflected successful learning (Figure 1b). Trials with licks at the tone onset were discarded from the analysis (Figure S1).

**Figure 1.**
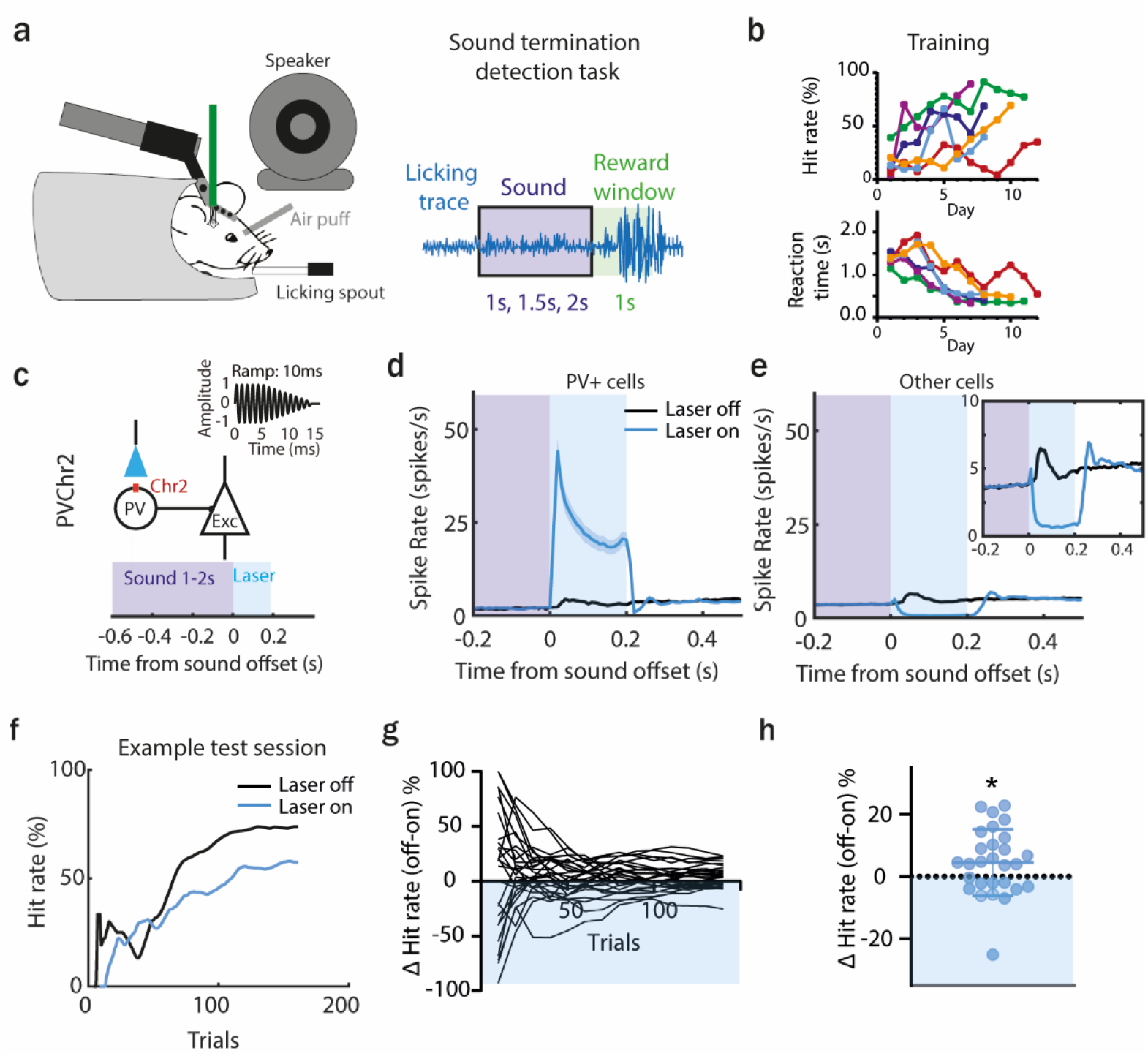
Preventing AAF offset responses decreases the ability of a mouse to detect sound termination. (**a**) Illustration of head-fixed behavioural set up. Piezo sensor attached to licking spout measured behavioural response to detection of sound termination (left). Schematic of the behaviour paradigm of offset detection task. Animal had to detect sound offset of 9 kHz PT played with three different durations: 1s, 1.5s, 2s within a reward window of 1s. (**b**) Indications of successful learning. Increasing hit rate of offset detection over training days (top, n=6 animals). Decreasing reaction time for offset detection over training days (bottom, n=6 animals). (**c**) Schematic of laser manipulation. Laser light was used for 200 ms following sound termination in animals expressing Chr2 in PV cells. Inset: zoom in. (**d**) Activity of PV+ cells (mean± SEM) following sound termination in laser on (blue) and laser off (black) trials, n=336 cells, 28 sessions, 6 animals. (**e**) Activity of other cells (mean± SEM) following sound termination in laser on (blue) and laser off (black) trials, n=2249 cells, 28 sessions, 6 animals. (**f**) Hit rates within an example sessions during offset detection task. Blue and black lines represent hit rate for laser on and laser off trials respectively. (**g**) Difference in hit rate between laser off and laser on trials during individual sessions (n=28 sessions, 6 animals) (**h**) Overall difference in correct hit rate for laser off and laser on trials at the end of the training (mean± SEM), p=0.0264, n=28 sessions, 6 animals, One sample Wilcoxon test. See also Figure S1.

After this training phase, animals underwent a craniotomy and the AAF was mapped. On the following day they were tested with tones played at 60 dB SPL. In half of the trials, a laser light was delivered above the AAF for 200 ms starting at sound termination (Figure 1c) in order to activate PV+ cells (Figure 1d) and to significantly reduce offset responses in non PV+ cells (Figure 1e). The comparison of the animals’ performance during laser on and laser off trials showed that preventing AAF offset responses significantly decreased the performance to detect sound termination (Figure 1f-h). This experiment demonstrates that offset responses in the AAF are behaviourally relevant.

### Bigger offset responses correlate with better detection of sound termination

Previous studies illustrated how offset responses in the ACx of rats and cats strongly depend on the fall time of a sound (Qin et al., 2007; Takahashi et al., 2004). We used faster fall-ramp to evoke higher amplitude offset responses and asked whether the amplitude of offset responses and the animal’s ability to detect sound termination are correlated.

We used a similar experimental paradigm as in Figure 1, where the animal had to detect the end of the 9 kHz tone played at 60 dB SPL with, this time, a fall-ramp of 10 or 0.01 ms. As expected from previous studies, fast fall–ramps (0.01 ms) lead to significantly higher offset responses than longer ones (10.0 ms), as tested during awake passive recordings (Figure 2a, Figure S2) or when animals were performing a behavioural task (Figure 2b). We confirmed that the sounds with a 0.01 ms fall-ramp are not causing an additional artificial onset response (Figure S3). The analysis of hit rates showed that mice could correctly detect when sounds ended for tones with both short and long ramps and no significant difference in detection rate between the two ramps were observed when looking at the end of test sessions (Figure 2c-e). However, at the beginning of the test sessions, mice were significantly better at detecting sound offset when the ramp was fast.

**Figure 2.**
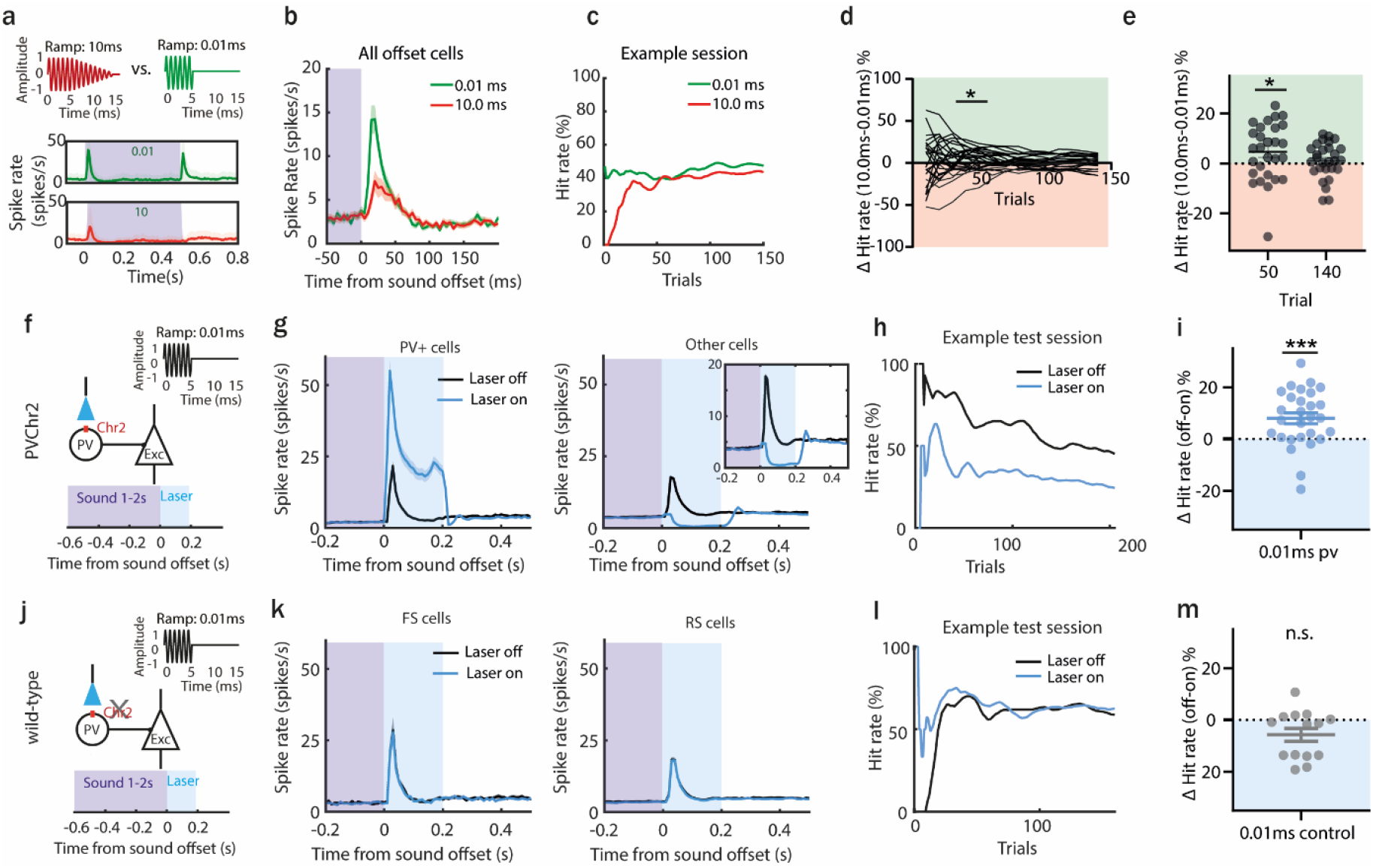
Bigger offset responses correlate with better detection of sound termination. (**a**) Schematic of sounds used in the behaviour task (0.01 ms or 10 ms fall-ramp) and PSTH (mean± SEM) of responses they evoked in acute AAF recordings. (**b**) PSTH (mean± SEM) averaged over AAF neurons (n=169, 2 animals) during sound termination detection task (green line: 0.01 ms offset ramp; red line: 10.0 ms offset ramp). (**c**) Hit rates of example session during offset detection task. Green and red lines represent hit rate for short (0.01 ms) and long (10 ms) offset ramps. (**d**) Difference in hit rate for sounds with short and long offset ramps through the behaviour session. Green and red shaded area indicate that offset detection was better for short or long ramps, respectively. (**e**) Comparison of correct hit rate for 0.01 ms and 10.0 ms ramp at Trial_50_ (p=0.0247, n=28 sessions, 10 animals, Wilcoxon test) and Trial_140_ (p=0.3052, n=28 sessions, 10 animals, Wilcoxon test). (**f**) Schematic of experimental design: sound of 9 kHz with 0.01 ms fall ramp was used and laser was applied for 200 ms following sound termination in animals expressing Chr2 in PV cells. (**g**) Activity of PV+ cells (mean± SEM) following sound termination in laser on (blue) and laser off (black) trials (left), n=336 cells, 28 sessions, 6 animals. Activity of other cells (mean± SEM) following sound termination in laser on (blue) and laser off (black) trials (right), n=2249 cells, 28 sessions, 6 animals. (**h**) Hit rates within an example sessions during offset detection task. Blue and black lines represent hit rate for laser on and laser off trials respectively. (**i**) Difference in correct hit rate for laser off and laser on trials at the end of the training (mean± SEM), p=0.0006, n=28 sessions, n=6 animals, One sample Wilcoxon test. (**j**) Schematic of control experiment: sound of 9 kHz with 0.01 ms fall ramp was used and laser was applied for 200 ms following sound termination in wild-type animals. (**k**) Population activity (mean± SEM) following sound termination in laser on (blue) and laser off (black) trials in fast (left), n=104 cells, 14 sessions, 3 animals, and regular spiking neurons (right), n=951 cells, 14 sessions, 3 animals. (**l**) Hit rates within an example sessions during offset detection task. Blue and black lines represent hit rate for laser on and laser off trials respectively. (**m**) Difference in correct hit rate for control animals in laser off and laser on trials at the end of the training (mean± SEM), p=0.0906, n=14 sessions, 3 animals, One sample Wilcoxon test. See also Figures S2-S4.

This suggests that sounds terminated with a fast fall-ramp, and therefore triggering a bigger offset response, were significantly easier to detect at the beginning of the sessions, when the task was still new and possibly more difficult than after exposure to more trials. This result is in line with previous findings showing cortical involvement in difficult but not easy tasks (Ceballo et al., 2019; Christensen et al., 2019; Dalmay et al., 2019; Kawai et al., 2015). There was no significant difference in the reaction times for both tested ramps (Figure S4).

To confirm that bigger offset responses help mice detecting sound termination, we performed another behavioural experiments with optogenetics, this time manipulating offset responses evoked by fast ramp (Figure 2f). As previously shown (Figure 1d, e), the laser significantly activated PV+ cells, resulting in the suppression of high amplitude offset responses in non-PV+ cells (Figure 2g). Minimizing high amplitude offset responses in the AAF significantly decreased the performance to detect sound termination (Figure 2h, i). To control that the light itself, without ChR2, was not causing any changes in behavioural performance, we repeated the same experiments in wild type animals (Figure 2j). The laser alone had no effect, neither at the neural (Figure 2k) nor at the behaviour level (Figure 2l, m). Together, these experiments confirm that changing offset responses in the AAF influences behavioural performance. They demonstrate that the neuronal activity following sound termination in the AAF is used by the animal to detect sound termination.

### The activity of AAF neurons in a sound termination detection task can be predictive of the animal’s performance

As the suppression of auditory offset responses in the AAF has an effect on performance, we asked if AAF activity during single trial could be predictive of the animal’s behaviour. We used a logistic regression model to predict the mouse’s action from the single-trial population activity (cross-validated, L2 penalty, see methods). We examined the classifier accuracy for the model trained and tested on *spontaneous, onset, sustained, offset* and *late* response (Figure 3a) from an equal number of hit and miss trials, from all experiments with fast ramp. We compared the classifier accuracy trained on different response type and found that offset and late responses allowed for significantly better action decoding than spontaneous or onset responses (Figure 3c). These results suggest that AAF offset responses can be informative on the animal’s decision. They emphasise again the behavioural relevance of AAF offset responses.

**Figure 3.**
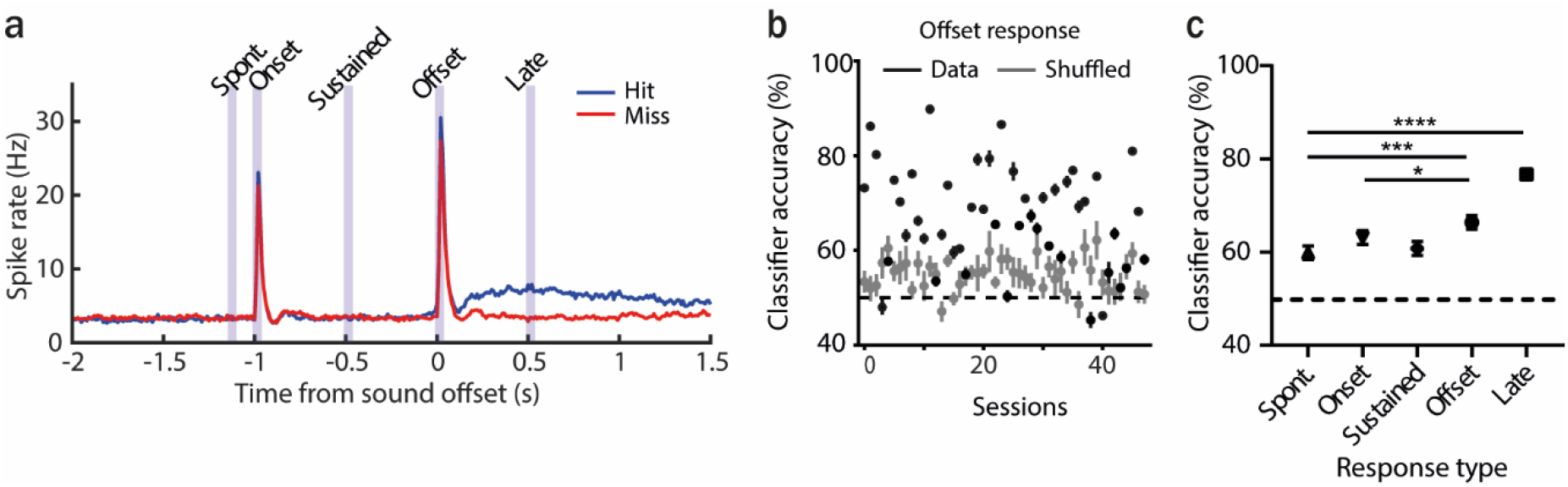
The activity of AAF neurons can be predictive of the animal performance. (**a**) Averaged PSTH (mean± SEM) of AAF neuron’s response to 1 s long PT (9 kHz) played at 60 dB SPL during hit (blue) and miss (red) trials, n=5076 cells, 59 sessions, 12 animals. (**b**) Classification accuracy based on offset responses (mean ± SEM) for real and shuffled data. (**c**) Comparison of classifier accuracy of decoders trained and tested on spontaneous activity, onset, sustained, offset and late response of AAF neurons (mean± SEM): spont. vs offset: p=0.0005; offset vs onset: p=0.0489; spont. vs late: p<0.0001, n=48 sessions, 12 animals, Wilcoxon test.

### The AAF is highly specialized for processing information on sound termination

Knowing that offset responses in the AAF are behaviourally relevant and influence sound termination perception, we next asked what mechanisms are driving these cortical offset responses and what properties distinguish them from subcortical ones.

We performed awake electrophysiological recordings in the AAF and the MGB and analysed the response profile dynamics of cells within both regions. We recorded multi-unit activity evoked by 50 ms pure tones (PT) with varying frequency (4 to 48.5 kHz) and sound level (0 to 80 dB SPL) presented with randomized inter-stimulus-intervals (ISI) (500–1000 ms).

K-means clustering of spike-sorted unit (SU) activity was used to identify cells with distinct temporal dynamics. The clustering method was performed on the averaged poststimulus time histogram (PSTH) in both MGB (n=779 SU, 5 animals) and AAF (n=346 SU, 6 animals) recordings pooled together. The analysis time window for the clustering was chosen to emphasize the offset rather than the onset responses (25 – 75 ms, bin size: 5ms). Davies Bouldin evaluation was used to determine the optimal number of clusters (Figure S5). Nine clusters were identified, reflecting five main temporal categories of auditory evoked responses: *onset-only, late onset, onset-offset, sustained* and *suppressed* (Figure 4). Few clusters with the same temporal dynamic pattern were detected (e.g. D-G) resulting from varied latencies, width and ratio of offset/onset responses. These clusters were merged for further analysis. In the MGB, both *onset-only* and *onset-offset* cells represented the biggest clusters: 40.1 % and 33.0 %, respectively (Figure 4b). Cells with these two temporal response patterns revealed also separate anatomical clusters within the MGB (Figure S6)(He, 2002). *Suppressed* responses were found in 12.3 %, *late onset* responses in 10.2 % and *sustained* in 4.4% of cells. In the AAF, most cells were *onset-offset* responsive (83.2 %). Other categories were represented by much smaller proportions: *onset-only* (3.2 %), *late onset* (6.1 %), and *sustained* (7.5 %). In contrast to the MGB, no *suppressed* cells were recorded in the AAF. The overrepresentation of onset-offset responses in the AAF compared to the preceding nucleus of the auditory pathway indicates that the AAF is highly specialized in processing information on sound termination.

**Figure 4.**
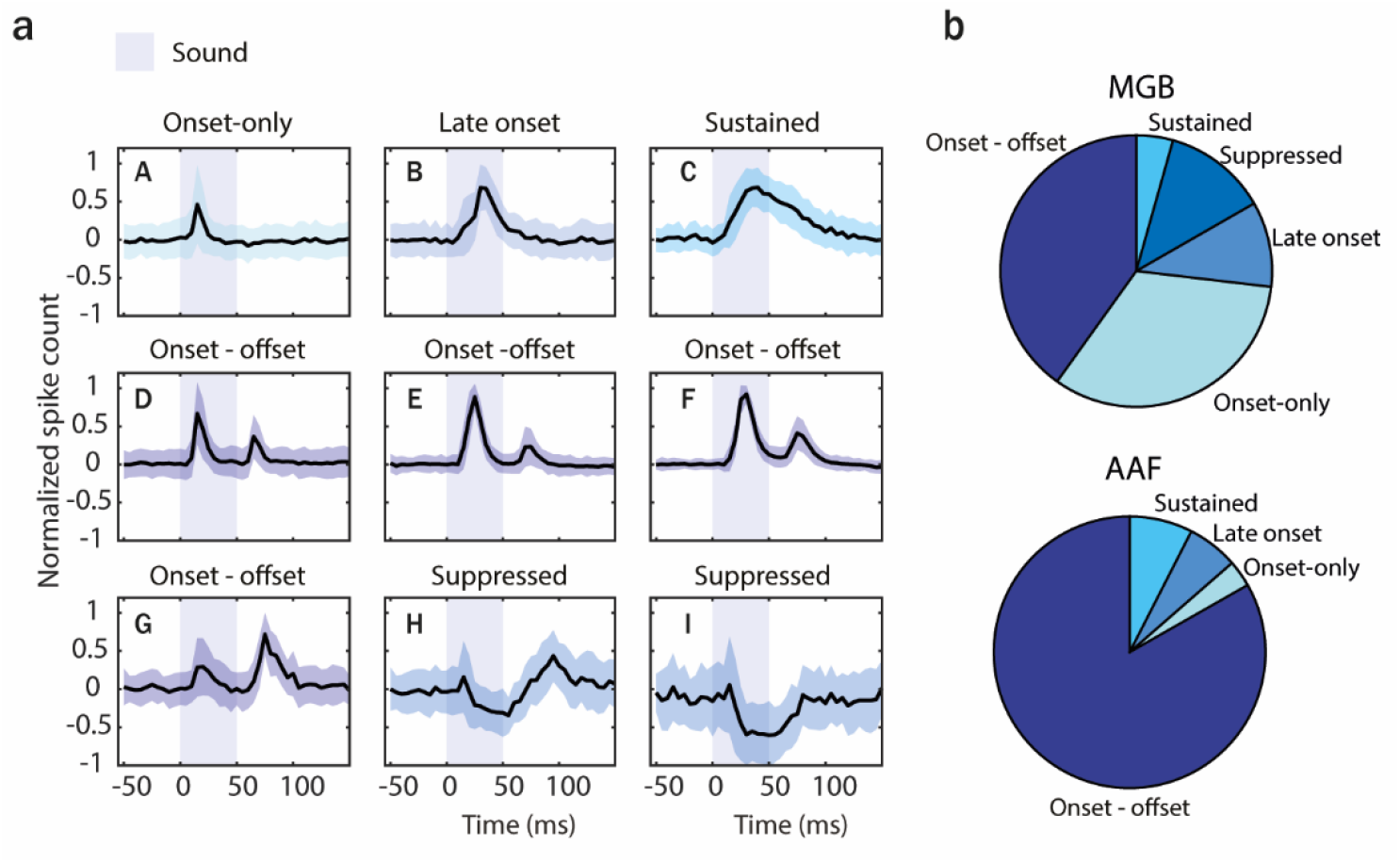
The AAF has significantly more offset-responsive neurons than its input nucleus. (**a**) Results of k-means clustering performed on both MGB (n=779) and AAF (n=346) neuron’s responses (time window: 25-75 ms, bin size: 5ms) evoked by 50 ms PT with varied frequency (4 to 48.5 kHz) and sound level (0 to 80 dB SPL) presented with randomized ISI (500–1000 ms). Graphs represent mean signal of cells belonging to each cluster. Data represent mean ± STD. The blue shaded bars represent the tone. (**b**) Representation of cells with distinct temporal dynamic of responses in the MGB and the AAF. See also Figures S5, S6.

### Onset-offset responsive cells are the main inputs from the MGB to the AAF

As the AAF contains cells with mainly onset-offset responses - unlike the MGB or A1 (Solyga and Barkat, 2019)-we asked whether these cortical offset responses were inherited from the MGB cells, or whether they arose *de novo* in the AAF. We combined *in vivo* electrophysiological recordings in the MGB (n=1548 SU, 5 animals) with antidromic stimulation of the AAF, followed by pure tone stimulation to characterize the temporal dynamic of cells projecting from the MGB to the AAF (Figure 5a, Figure S7). A stimulating electrode was inserted into the previously functionally identified AAF (see methods).

**Figure 5.**
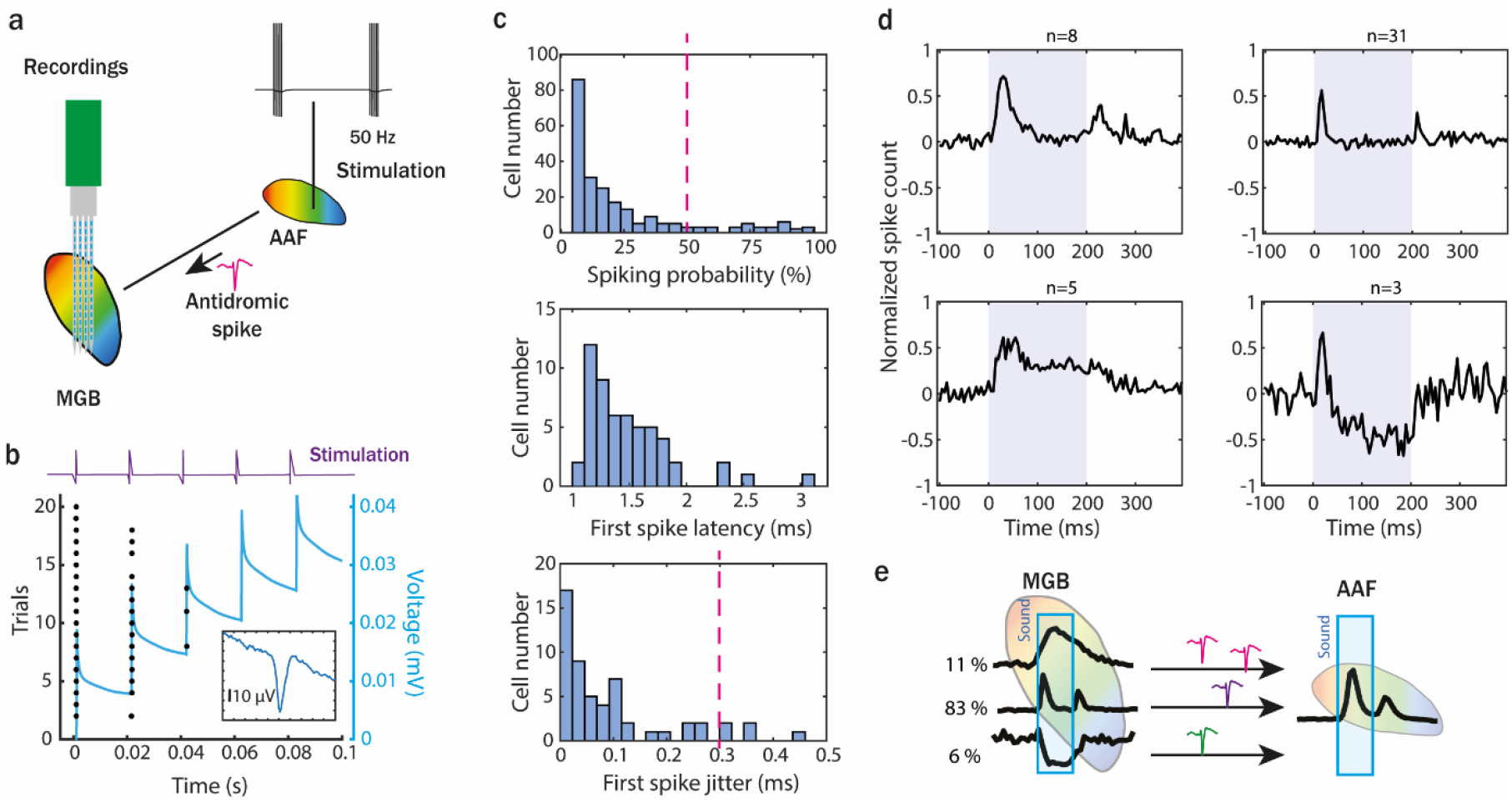
Onset-offset responsive cells are the main inputs from the MGB to the AAF. (**a**) Illustration of experimental set up to perform an antidromic stimulation of MGB neurons projecting to the AAF. Stimulation tip was placed in the AAF after field identification (based on functional tonotopy). Monophasic electric pulses were delivered with 50 Hz at 30 µA. 64-channels electrode was inserted into MGB to record antidromic activity. (**b**) Example MBG unit spike raster of antidromic activity within 20 trials (5 pulses in each trial). Blue line represents increasing current injected during electric stimulation. (**c**) Number of recorded antidromic spikes in MGB neurons, where at least 1 spike following stimulation was detected in time window from 1 to 5 ms after stimulation (*top*). Latency (*middle*) and jitter (*bottom*) of first antidromic spikes in MGB cells which were detected in at least half of the electric stimulation trials. (**d**) Temporal dynamic of clustered MGB cells (k-means) projecting to AAF identified during antidromic experiment (mean± SEM). Three main temporal categories of auditory evoked responses were identified: onset-offset, sustained and suppressed. MGB cells were considered as AAF input if (1) in more than 50% of trials antidromic spikes were detected and (2) antidromic spikes jitter was lower than 0.3 ms. (**e**) Illustration of temporal dynamic and proportion of MGB cells projecting to AAF based on our antidromic results. See also Figure S7.

Pulse trains of monophasic square pulses were used for the electric stimulation (Figure 5b). We identified the MGB cells directly connected to the AAF by analysing the percentage and latency of responses to the first stimulation pulse in each train. The MGB cells that fired as a response to the AAF stimulation in at least 50% of the trials with a first spike latency of 1 to 3 ms and a trial-to-trial latency jitter lower than 0.3 ms (Serkov, 1976) were considered to be sending direct inputs to the AAF (Figure 5c). These MGB cells were clustered mainly as onset-offset cells (83 %). We also identified some sustained (11 %) and suppressed (6 %) cells projecting from the MGB to the AAF, but their representation was significantly lower. Onset-only cells were not identified (Figure 5d). These results indicate that offset responses in the AAF are mainly inherited from the MGB. Whether further processing of these offset responses within the cortex took place is however still unclear.

### Offset responses increase with sound duration mainly in the AAF

Given the presence of offset responses in the MGB and the AAF, we next asked whether their properties were similar in both regions. To reveal differences in offset processing we decided to check how offset responses in the MGB and the AAF are affected by different sound properties such as sound duration, spectral content or temporal complexity. We first addressed the question of the dependence of offset responses on sound duration (Scholl et al., 2010; Solyga and Barkat, 2019). We recorded responses in the MGB and the AAF offset cells (clustering based on Figure 4) to 60 dB SPL tones with durations varying between 50 and 500 ms and inter-stimulus intervals varying between 50 and 2000 ms (Figure 6a, d). For the MGB, the tone frequency was dependent on the offset BF of neurons in each session (because of narrow offset tuning of MGB neurons); for the AAF, a fixed PT of 9 kHz was used (as most of the widely tuned cells were responding to this frequency).

**Figure 6.**
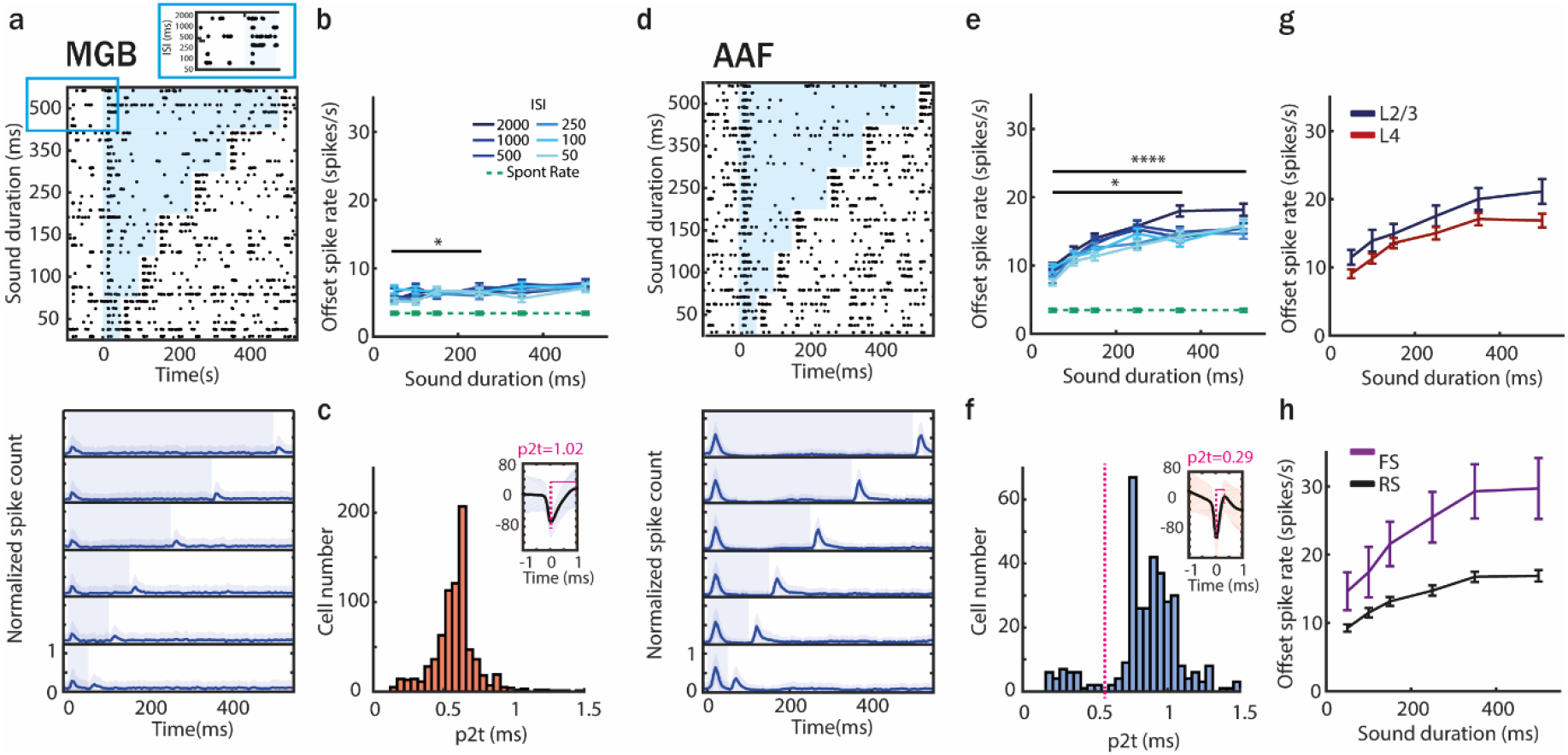
Offset responses increase with sound duration mainly in the AAF. (**a**) Raster plot of an example MGB neuron’s response to PT (frequency chosen based of offset tuning) with sound duration varying between 50 and 500 ms and ISI between 50 and 2000 ms (*top*) and PSTH (mean± STD) averaged over all neurons population (*bottom*). The blue shaded bars represent the tone. (**b**) MGB neurons offset responses (mean± SEM) to PT with increasing duration across all ISI of 2000 ms (correlation between sound duration and response rate: PT, ρ=0.05, p=0.02, n=307 SU, 5 animals, Spearman correlation). Comparison of offset responses: 50 ms vs 100 ms: n.s. p=0.08; 50 ms vs 150 ms: n.s. p=1; 50 ms vs 250 ms: p=0.0318; 50 ms vs 350: n.s. p=0.7013; 50 ms vs 500 ms n.s. p=0.9064, Dunn’s multiple comparisons test (**c**) Distribution of peak-to-trough times (p2t) of MGB neurons. (**d**) Raster plot of an example AAF neuron’s response to PT (9 kHz) played at 60 dB SPL with sound duration varying between 50 and 500 ms and ISI between 500 and 2000 ms (*top*) and PSTH (mean± STD) averaged over all neurons population (*bottom*). The blue shaded bars represent the tone. (**e**) AAF neurons offset responses (mean± SEM) to PT with increasing duration across ISI of 2000 ms (correlation between sound duration and response rate: PT, ρ=0.25, p<0.0001, n=285 SU, 6 animals, Spearman correlation). Comparison of offset responses: 50 ms vs 100 ms: n.s. p=0.5655; 50 ms vs 150 ms: n.s. p=0.9983; 50 ms vs 250 ms n.s. 0.0819; 50 ms vs 350: p<0.0411; 50 ms vs 500 ms p<0.0001, Dunn’s multiple comparisons test. (**f**) Distribution of p2t of AAF neurons (**g**) Comparison of offset spike rate in L2/3 and L4 neurons in AAF for sounds with duration varying between 50 and 500 ms and longest tested ISI of 2000 ms (mean± SEM), L2/3: ρ=0.26, p<0.0001 n_2/3_=87, L4: ρ=0.24, p<0.0001 n_4_=198, Spearman correlation. (**h**) Comparison of offset spike rate of fast and regular spiking neurons in AAF for sounds with duration varying between 50 and 500 ms and longest tested ISI of 2000 ms (mean± SEM), FS: ρ=0.29, p=0.00012, n_FS_=28, RS: ρ=0.24, p<0.0001, n_RS_=257, Spearman correlation.

The correlation between sound duration and offset spike rate evoked by tones in onset-offset cells was very weak in the MGB but much stronger in the AAF (Figure 6b, e). In the MGB, population responses showed almost no difference in offset spike rate evoked by tones when durations changed between 50 ms and 500 ms (Figure 6b). In the AAF however, differences in tone duration were significantly reflected by increased offset spike rates for the longest sounds (Figure 6e).

To explore whether the dependence of offset responses on sound duration is a result of AAF computations in layer 2/3 (L2/3) or is already present in the input layer 4 (L4), we compared the dependence of offset responses on sound duration in these layers (Figure 6g). Our recordings span the range of 150 to 600 µm from the pia surface, corresponding mainly to L2/3 (150-300 µm) and L4 (300-500 µm) (Meng et al., 2017). A significant correlation was present both in L2/3 and L4, suggesting that the increase in offset response amplitude with sound duration is not unique to one layer.

As the MGB does not contain fast spiking interneurons (FS) (Bartlett, 2013), we then asked whether their presence in the AAF could be driving the dependence of offset responses on sound duration in this cortical region (Figure 6c, f). We distinguished putative FS and regular spiking (RS) neurons based on the peak-to-trough times (p2t) of their spike waveforms (Figure 6f). Fast spiking units were defined as having a p2t smaller than the minimum between the two peaks of the p2t distribution (0.55 ms), in accordance with previous studies (Moore and Wehr, 2013). The unimodal distribution of p2t in the MGB confirmed the lack of FS in this region (Figure 6c). We found a significant correlation between offset spike rate and sound duration both in FS and RS (Figure 6h) neurons, ruling out the possibility that one of these cell populations is alone driving the dependence of offset responses on sound duration in the AAF.

Together, these results reveal an important difference between AAF and MGB offset encoding and demonstrate a clear amplification of the dependence of offset responses on sound duration in the AAF as compared to the MGB.

### Offset responses to white noise stimulation are present in the AAF but not in the MGB

Next, we compared offset responses in the MGB and the AAF evoked by sounds with different spectral complexity. We recorded responses in both regions to 500 ms pure tones (PT) and white noise (WN) bursts played at 60 dB SPL (for the MGB the PT frequency was chosen based on offset BF of neurons in each session; for the AAF, a fixed PT of 9 kHz was used). Our results showed very distinct neuronal activity patterns as a response to WN or the spectrally less complex PT, both in the MGB and in the AAF (Figure 7). In the MGB, 500 ms WN actually evoked no offset responses above spontaneous activity, unlike PT of the same length (Figure 7a, c). In the AAF however, both PT and WN did evoke offset responses (Figure 7b, d). The lack of offset responses evoked by WN in the MGB, and their significant presence in the AAF, reveals offset responses generated *de novo* in the cortex.

**Figure 7.**
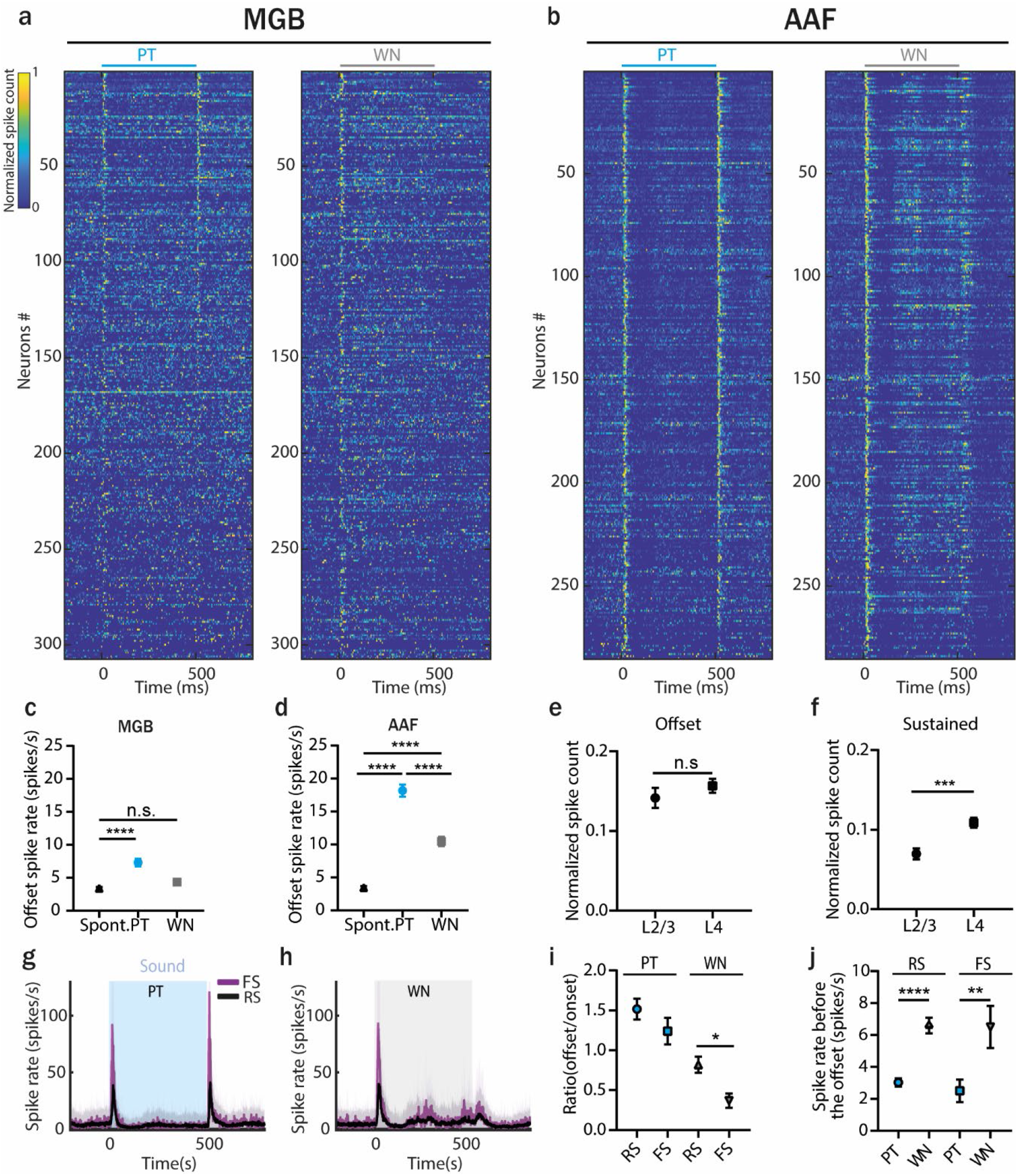
Offset responses to white noise stimulation are present in the AAF but not in the MGB. (**a**) Normalized PSTH of MGB neurons to 500 ms PT or WN bursts, bin size: 5ms. Data are sorted by descending spike rate at the PT offset. (**b**) Normalized PSTH of AAF neurons to 500 ms PT or WN bursts, bin size: 5ms. Data are sorted by descending spike rate at the PT offset. (**c**) Comparison of MGB offset responses evoked by PT and WN for onset-offset cells. Data represent mean ± SEM, PT vs spont. rate: p<0.0001; WN vs spont. rate: p=0.071, n=307, Wilcoxon test. (**d**) Comparison of AAF offset responses evoked by PT and WN for onset-offset cells. Data represent mean ± SEM, PT vs spont. rate: p<0.0001; WN vs spont. rate: p<0.0001, PT vs WN: p<0.0001, n=285 Wilcoxon test. (**e**) Comparison of sustained responses (calculated in window: 100-500ms) for AAF cells from L2/3 and L4 evoked by 500 ms WN stimulation (mean± SEM), p=0.0003, n_2/3_=87, n_4_=198, Mann-Whitney test. (**f**) Comparison of offset responses for AAF cells from L2/3 and L4 evoked by 500 ms WN stimulation (mean± SEM), p=0.2963, n_2/3_=87, n_4_=198, Mann-Whitney test. (**g, h**) Averaged PSTH of fast (n=29) and regular (n=249) spiking AAF neuron’s response to PT and WN bursts played at 60 dB SPL with sound duration 500 ms and ISI between 500 and 2000 ms. (**i**) Ratio of offset/onset responses evoked by 500 ms PT or WN in fast and regular spiking AAF neurons (mean± SEM), p=0.0202, n_FS_=28, n_RS_=246, Mann-Whitney test. (**j**) Spike rate preceding sound offset (calculated in window: 450-500ms) in AAF neurons for longest ISI of 2000 ms (mean± SEM), RS: p<0.0001, n=257; FS: p=0.0013, n=28 Wilcoxon test. See also Figure S8.

We then asked whether offset responses to WN stimulation differed between different neuronal populations of the AAF. We found that responses to WN were significantly more sustained in L4 than in L2/3 (Figure 7e, calculated in a window of 100-500 ms following sound onset), despite having similar offset responses in both layers (Figure 7f). It seems that responses to WN stimulation, even if not present among MGB inputs, arise already in the AAF input layer. Next, we compared offset responses evoked by PT and WN in putative FS and RS neurons (Figure 7g, h). As expected, offset responses were bigger and faster in FS than RS neurons following PT termination (Figure 7I, Figure S8a, b). In addition, there was not a significant difference in spike rate and latency between FS and RS neurons following WN termination (Figure S8c). The comparison of ratios of offset/onset responses between cell and sound types showed that the ratio for FS neurons was significantly smaller than the ratio for regular spiking neurons with WN stimulation (Figure 7i). This relative decrease of offset responses in FS neurons could reduce inhibition and result in an enhanced activity of excitatory cells, leading to offset responses generated *de novo* in the AAF. These findings suggest that FS neurons could play an important role in cortical processing of WN offset responses.

The PSTHs of AAF neurons responding to PT and WN stimulation indicate that WN evokes more sustained activity than PT (Figure 7b). More specifically, WN gives rise to sharp onset responses followed by suppression, and then an activity rebound at around 200 ms followed by a second suppression phase and another rebound. Could this specific response of neurons to WN stimulation influence the strength of AAF offset responses? Comparing firing rate of FS and RS cells in the AAF 50 ms before the sound termination revealed that they were significantly bigger for WN than PT stimulation (Figure 7j). This extended firing could possibly affect the generation of offset for WN stimulation.

Studying offset responses evoked by WN bursts revealed two very interesting differences between MGB and AAF processing. First, offset responses to WN stimulation are not present in the MGB but are present in the AAF. Second, cells in the AAF seem to follow ongoing WN stimulation with some kind of bursting activity happening every ∼200 ms.

### Offset responses encode more than just silence

The precise detection of fast changes in sound frequency and level is crucial for gap detection and vocalization (Kopp-Scheinpflug et al., 2018; Sollini et al., 2018). We asked if offset responses in the MGB and the AAF could encode more than silence - i.e. sound termination - such as important changes within temporally discontinuous sounds (Lu et al., 2001). We recorded responses to a multi-frequency component sound in the MGB (n=275 SU, 5 animals) and in the AAF (n=284 SU, 6 animals). The complex sound consisted of three frequency components (20 kHz, 14 kHz and 9 kHz) played at 60 dB SPL, which had a common onset but ended at different time points (300 ms, 400 ms, 500 ms). The offset responses evoked by removing one or two frequency components demonstrate that neurons can encode the disappearance of a frequency component in an ongoing sound, especially in the AAF (Figure 8a-f).

**Figure 8.**
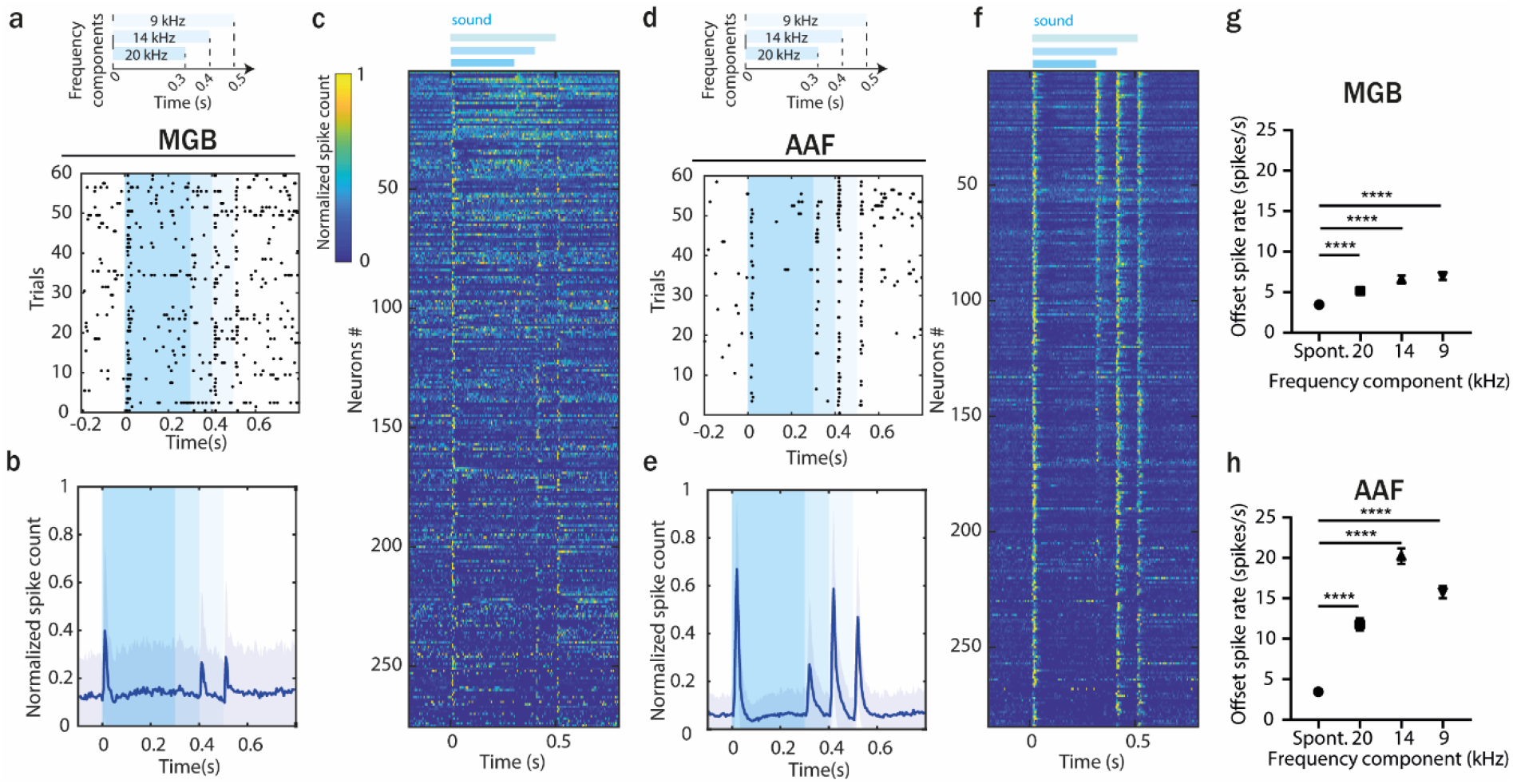
Offset responses encode more than just silence. (**a**) Raster plot of an example MGB neuron’s response to 3-components stimuli. (**b**) Averaged PSTH (mean ± STD) of MGB neuron’s response to 3-components stimuli, n=275, n=5 animals (**c**) Normalized PSTH of MGB neuron’s response to 3-components stimuli, bin size: 5ms. Data are sorted by descending spike rate at the first component termination. (**d**) Raster plot of an example AAF neuron’s response to 3-components stimuli. (**e**) Averaged PSTH (mean ± STD) of AAF neuron’s response to 3-components stimuli, n=284, n=6 animals. (**f**) Normalized PSTH of AAF neuron’s response to 3-components stimuli, bin size: 5ms. (**g**) Comparison of spiking rate of MGB neurons following removal of each frequency component. Data represent mean ± SEM, p<0.0001, n=275, n=5 animals Wilcoxon test. (**h**) Comparison of spiking rate of AAF neurons following removal of each frequency component. Data represent mean ± SEM, p<0.0001, n=284, n=6 animals Wilcoxon test.

A single MGB neuron usually encoded the removal of one or two frequency components. In contrast, most AAF neurons encoded the removal of all frequency components. However, at the population level the termination of all three components was significantly encoded in both MGB and AAF activities (Figure 8g, h). Interestingly, removing the first frequency component (20 kHz) evoked the smallest offset response, which, in the MGB, was close to spontaneous activity. Removal of the last component (9 kHz), which was followed by silence, evoked the highest offset response within MGB neurons. In contrast, the highest offset response was present in the AAF after removal of the second component (14 kHz). AAF neurons seem to have a stronger ability to integrate information over spectrally and temporally complex ongoing sounds and not only to respond to simply silence. This ability to encode important changes within ongoing sound could be crucial for processing temporally discontinuous sounds, which makes the AAF an interesting field to study the encoding of vocal calls.

## DISCUSSION

As the auditory system very robustly represents timing information, it is a model of choice to study neural offset responses evoked by the disappearance of a stimulus. In this study, we show that minimizing AAF offset responses decreases the mouse performance to detect sound termination, thus revealing their importance at the behavioural level. By combining *in vivo* electrophysiology recordings in the AAF and the MGB of awake mice, we also demonstrate that the AAF inherits, amplifies and sometimes even generates *de novo* offset responses. These results are of high importance for all studies on sensory processing, as the mechanisms determining specificities in cortical versus thalamic processing revealed by our studies could be common between the different sensory areas.

The functional significance of offset responses was long under debate (Saha et al., 2017). Here we show that minimizing AAF offset responses significantly decreases the performance of mice to detect sound termination (Figure 1, Figure 2). In addition, sounds terminating with a fast fall ramp and therefore triggering a bigger offset response, are significantly easier to detect at the beginning of a behavioural session when the task is still new and more difficult, than after exposure to more trials. A possible explanation for this dependence on task difficulty could be related to the distinct involvement of the ACx during more or less challenging tasks. Previous studies have indeed shown that the cortex could be required for difficult tasks, but less so for easier tasks or when the task is learned well (Ceballo et al., 2019; Christensen et al., 2019; Dalmay et al., 2019; Kawai et al., 2015). When the task is easy or familiar, high and low amplitude offset responses seem to be informative about sound termination to the similar extent. Finally, we used logistic regression to show that AAF activity during single trial could be predictive of the animal’s performance (Figure 3). The neural activity during hit trials in the AAF could be influenced by the animal’s general motivation (Fritz et al., 2003), motor related inputs (Schneider, 2020) or reward expectations (De Franceschi and Barkat, 2020).

Another approach to understand the behavioural role of auditory offset responses could be to study where the AAF is projecting and what could be the use of offset responses in these areas. It was previously shown that offset responses in the secondary auditory cortex are plastic and able to enhance the representation of a newly acquired, behaviourally relevant sound category (Chong et al., 2020). Whether the activity of AAF neurons is crucial for this plasticity to happen remains to be elucidated. Only recently, Nakata et al. revealed direct AAF connections to the secondary motor cortex, the primary somatosensory cortex, the insular auditory cortex and the posterior parietal cortex (Nakata et al., 2019). What the role of auditory offset responses in these fields could be and whether they provide any information for association with the somato-motor system remain unanswered.

Spectro-temporal tuning properties of auditory neurons differ during the presentation of natural and synthetic stimuli (David et al., 2007; Theunissen et al., 2001). Natural sounds also commonly start in an abrupt way, but their termination is obscured by sound reverberations, and therefore stops less sharply than synthetic stimuli. In many of our experiments, we used a sharp 0.01 ms sinusoidal offset ramp, while keeping the onset ramp at 4 ms. As a sharp ramp might cause a spectral splatter, we decided to quantify it in our protocol. Using an ultrasensitive microphone, we recorded acceleration traces of 9 kHz pure tone played at 60 dB SPL with 0.01 ms and 1 ms rise and fall-ramp (Figure S3a, b). For the fast fall ramp, we detected a weak spectral splatter present for less than 0.5 ms and covering the frequency range between 6 and 30 kHz (Figure S3c, d). We then investigated if this spectral splatter could trigger any significant onset response that would be mixed with the real offset responses. First, we asked if neurons with a BF within the spectral splatter frequency range displayed a bigger response to sound termination. We did not see any increased offset response in those neurons as compared to neurons with other BF (Figure S3e), indicating that putative onset responses to the splatter cannot be significant. Then, we asked if offset responses evoked by fast and slow ramps correlate. If fast offset responses were rather onset responses to the spectral splatter instead of real offset responses, no correlation between these onset responses and the offset responses triggered by a slow ramp would be expected. We showed that the offset responses to fast and slow ramps indeed highly correlate (Figure S3f, ρ=0.71, p<0.0001, Spearman correlation), suggesting that our protocol allows us to identify real offset responses. Finally, we asked if the responses at sound termination would influence the onset responses of a tone played shortly after. As offset and onset responses are driven by different sets of synapses (Scholl et al., 2010), we would expect the onset responses of the second sound to be affected by the first sound only if the response at sound termination were onset responses to the artefact, but not if they were real offset responses. We found that onset responses are suppressed by preceding onset responses but not affected by preceding offset responses (Figure S3g), even when interval was as short as 50 ms. This suggests that the offset responses we recorded are driven by different sets of synapses that onset responses, and can therefore not be onset responses. Together, these quantifications confirm that the neuronal responses measured at sound termination are not affected by the weak and short spectral splatter caused by a fast offset ramp. In the future, the role of offset responses in detecting sound termination should be studied in natural environments, using for example vocalization calls (Chong et al., 2020). This would tell if the auditory system, and the MGB or the AAF more specifically, evolved in a way to meet the challenges of detecting naturally terminating sounds.

Our results demonstrate that cortical offset responses are not only inherited from the periphery (Figure 4, Figure 5) but also amplified by sounds with longer duration (Figure 6). Could the presence of FS interneurons in the AAF, but not in the MGB (Bartlett, 2013), be driving the dependence of offset responses on sound duration in this cortical region? The strength of the correlation between offset responses and sound duration was similar between FS and RS cells in the AAF (Figure 6). However, FS neurons showed significantly bigger amplitude of offset responses in comparison to RS neurons. The AAF was previously shown to have more PV+ cells than the other auditory primary region A1 (Reinhard et al., 2019). Mechanistically, we suggest that such a prominent PV network in the AAF and the big amplitude offset responses they exhibit could be crucial for evoking duration dependent offset responses in the AAF but not in the A1 (Solyga and Barkat, 2019) or the MGB (Figure 6). With our antidromic experiments, we also show that some of the AAF cells receive inputs from sustained and suppressed MGB cells (Figure 5). Whether or not the inputs from these cells are involved in the duration dependence of offset response in the AAF should be explored further.

What could be the role of cortex in tracking subtle differences in sound duration? At the behavioural level, one could speculate that the amplitude of offset responses would be needed to better track subtle differences in sound duration, especially for sounds shorter than 500 ms, covering a spectrum of most mouse calls (Geissler and Ehret, 2002). Nevertheless, if the increase in amplitude of offset responses with sound duration is really a carrier of useful information or just a result of cortical cells being unable to handle short sounds remains unclear.

The spectral complexity of sounds is significantly modulating offset responses in the central auditory system in several ways. First, the lack of offset responses to WN in the MGB onset-offset cells and their presence in the AAF reveals offset responses generated *de novo* in the cortex (Figure 7). How could the differences in offset responses to WN stimulation be explained? The large spectral integration of thalamic inputs (Figure S9) in individual cortical neurons (Liu et al., 2007; Vasquez-Lopez et al., 2017) should not play a role, as no firing upon WN termination was observed in the thalamus. Interestingly, offset responses to WN stimulation were present in the AAF both in L4 and L2/3, suggesting that it is a general property of the AAF network to generate offsets *de novo* and not only an exclusive property of superficial layer. What mechanisms could drive offset responses to WN in the AAF? FS and RS neurons exhibit similar offsets responses to WN stimulation, thus not allowing us to speculate on their special involvement in *de novo* generated offset responses. The potential role played by other types of cortical neurons, like the somatostatin interneurons previously suggested to be involved in offset response generation (Liu et al., 2019), remains to be elucidated. It is also possible that information on sound termination arises from non-lemniscal areas projecting to the AAF. Such possible projections and their potential contributions have not been described yet.

Second, the activity during WN stimulation in the AAF shows a clear pattern of bursting activity, where a transient onset is followed by a suppression phase, then rebound activity around 200 ms, followed by another suppression and rebound phase. Does this reflect an internal clock allowing the AAF to follow sound duration irrespectively from inputs coming from the MGB? What could be the role of such bursting activity in the AAF and how could it be generated? Bursts are thought to be emitted by many subcortical and cortical areas of the brain, but their hypothesized functions differ across brain areas (Zeldenrust et al., 2018). It has previously been shown in marmosets that the auditory cortex can use a combination of temporal and rate representation to encode the wide range of complex, time-varying sounds (Lu et al., 2001). The offset responses and bursting activity we see in the mouse AAF could play these multiple roles in encoding different temporal features such as ongoing sound and its termination. If the bursting and offset activity observed in the AAF is a common feature of other primary sensory cortices is unclear.

Finally, we demonstrate that offset responses to WN or PT are strikingly different in the central auditory system (Figure 7). This could have multiple origins. First, the significantly increased activity of AAF neurons preceding WN termination which could in turn result in a decreased ability of neurons to respond properly to the end of the sound (Figure 7j). Second, the lack of proper inhibition - excitation balance (Figure 7i) could decrease the offset responses evoked by WN burst. WN stimulation is widely used in auditory research, especially for gap in noise detection (Syka et al., 2002; Threlkeld et al., 2008; Weible et al., 2014a; Weible et al., 2014b) and for offset responses studies (Anderson and Linden, 2016). It is an attractive auditory stimulus to study purely temporal information as it ensures an effective stimulation of the auditory system irrespectively of neuronal tuning. However, one has to be careful about generalizing results obtained with this stimulus to all auditory inputs, as our observations indicate that different mechanisms might be at play when WN or PT, and by extension natural sounds, are heard.

The experiments with multi-frequency component sounds show that within both the MGB and the AAF, offset responses indicate not only when a sound ends, but also all important changes that occur within a temporally discontinuous sound (Figure 8), emphasising their possible relevance for behaviour and perception. The temporal integration of offset responses is crucial for perceptual grouping of communication sounds, in which rapid changes in intensity and frequency occur (Sollini et al., 2018). Our results suggest that this integration is accentuated in the cortex, making it an interesting hub to look for the mechanisms that might explain impairments in sound-offset sensitivity, and by extension, deficits in temporal processing arising both in aging and disease. A deeper knowledge of the cellular and circuit mechanisms of cortical offset responses could be crucial to develop new strategies to prevent abnormal auditory perceptual grouping.

## MATERIALS and METHODS

### Surgical procedures

All experimental procedures were carried out in accordance to Basel University animal care and use guidelines, and were approved by the Veterinary office of the Canton Basel-Stadt, Switzerland. 35 C57BL/6J mice were used in this study, procured from Janvier LABS, France. Mice were a mixture of males and females and aged between 7 and 12 weeks of age at the time of behavioural training or electrophysiological recording.

Awake electrophysiology recordings and behaviour experiments were performed on adult (7–12 weeks) male and female C57BL/6J mice (Janvier, France). For surgeries mice were anesthetized with isoflurane (4% for induction, 1.5 to 2.5% for maintenance) and subcutaneous injection of bupivacain/lidocain (0.01 mg/animal and 0.04 mg/animal, respectively) was used for analgesia. A custom-made metal headpost was fixed with super glue (Henkel, Loctite) on the bone on top of the left hemisphere, and used to head-fix the animals. Their body temperature was kept at 37°C with a heating pad (FHC, ME, USA) and lubricant ophthalmic ointment was applied on both eyes. Craniotomy (∼2×2 mm^2^) was performed with a scalpel just above the right auditory cortex and covered with silicone oil and silicone casting compound (Kwik cast, World Precision Instruments, Inc. FL, USA) during the 2 h recovery period from the anesthesia.

### Recordings

The electrophysiological recordings were performed in awake mice (AAF: n=6; MGB: n=5). Mice were head fixed and placed in the cardboard tube for recordings inside a sound box. Extracellular recordings were conducted in AAF (identified based on the functional tonotopy: ventro-dorsal increase in BF) and MGB (centered 0.8 mm anterior and 2 mm lateral to Lambda). Multi-channel extracellular electrodes with 32 channels (A4×8-5 mm-50-200-177-A32 or A1×32-5mm-25-177-A32 Neuronexus, MI, USA) or 64 channels (A4×16-5mm-50-200-177-A64, Neuronexus, MI, USA) were inserted orthogonal to the brain surface with a motorized stereotaxic micromanipulator (DMA-1511, Narishige, Japan) at a constant depth (AAF: tip of electrode at 556±9 µm from pia; MGB: tip of electrode at 3575±300 µm from pia). Responses from extracellular recordings were digitized with a 32- or 64-channel recording system (RZ5 Bioamp processor, Tucker Davis Technologies, FL, USA) at 24414 Hz. Single unit clusters were identified from raw voltage traces using kilosort (Pachitariu et al., 2016) (CortexLab, UCL, London, England) followed by manual corrections based on the inter-spike-interval histogram and the consistency of the spike waveform (phy, CortexLab, UCL, London, England) and further analysed in MATLAB (Mathworks, MA, USA).

### Auditory stimulation

Sounds were generated with a digital signal processor (RZ6, Tucker Davis Technologies, FL, USA) at 200 kHz sampling rate and played through a calibrated MF1 speaker (Tucker Davis Technologies. FL, USA) positioned at 10 cm from the mouse’s left ear. Stimuli were calibrated with a wide-band ultrasonic acoustic sensor (Model 378C01, PCB Piezotronics, NY, USA).

### Antidromic stimulation

To identify temporal dynamics of cells projecting from MGB to AAF *in vivo* electrophysiology recordings in MGB were combined with antidromic stimulation of AAF (n=5 animals). First Mice were anesthetized with intraperitoneal injection of ketamine/xylazine (80 mg/kg and 16 mg/kg, respectively) and subcutaneous injection of bupivacain/lidocain (0.01 mg/animal and 0.04 mg/animal, respectively) was used for analgesia. Ketamine (45 mg/kg) was supplemented during surgery as needed. For surgery, mice were head fixed and their body temperature was kept at 37 °C with a heating pad (FHC, ME, USA). Two separate craniotomies (∼2 × 2 mm^2^) were performed with a scalpel above the right MGB and auditory cortex and covered with silicone oil. AAF was mapped with electrophysiology recordings based on ventro-dorsal increase in BF to identify target area for stimulus pipette insertion. Then both craniotomies were covered with silicone casting compound (Kwik cast, World Precision Instruments, Inc. FL, USA) during the 2 h recovery period from the anaesthesia. Electric stimulator (Master-8, A.M.P.I., Israel) was connected to a stimulation isolator (ISO-Flex, A.M.P.I, Israel), which was then connected to the wire electrode. Wire electrode was fixed in pulled capillary glass (tip:<10 µm) filled with saline and then inserted into AAF (∼300 µm). Monophasic square wave pulse were generated with electronic stimulator as pulse train (pulse duration: 0.1 ms; frequency: 50 Hz; train number: 20; intensity: 30 uA (similar to method described in (Peng et al., 2017)). In the same time electrophysiology recordings with 64-channels electrode were performed in MGB. As described in recordings section, spike sorting was performed using kilosort (Pachitariu et al., 2016), followed by manual corrections in phy, and further analysis in MATLAB. To ensure the absence of electrical artefacts, mean cluster waveform from raw data was calculated for each antidromicaly-identified cluster. Clusters containing any high amplitude electric artefacts were removed from the analysis.

### Offset detection task

#### Headplate implant

Mice were implanted with a custom-made metal headpost at 7-8 weeks after birth under isoflurane anesthesia (4% induction, 1.2 to 2.5% maintenance). Local analgesia was provided with subcutaneous injection of bupivacaine/lidocaine (0.01 mg/animal and 0.04mg/animal, respectively). A headpost and a ground screw were fixed to the skull with dental cement (Super-Bond C&B; Sun Medical, Shiga, Japan). The portion of skull above the target recording site was left free from cement, and covered with a thick layer of Kwik-Cast Sealant (WPI, Sarasota, FL, USA). Post-operative analgesia was provided with an intraperitoneal injection of buprenorphium (0.1 mg/kg). After recovery from the surgery for a couple of days, mice were food restricted. *Training*. Mice were then placed in the cardboard tube and adapted to the head restraint. The speaker was placed 10 cm away from the left ear of the animals. Next they were taught to associate a sound offset with a reward availability. Mice were trained to detect sound offset of pure tones (9 kHz) played at 80 dB SPL (training) with varied duration (1 s, 1.5 s, 2 s). Rise ramp of the tones was always fixed to 10.0 ms, while at the offset fast (0.01 ms) or slow (10.0 ms) ramp was used and varied randomly. During the beginning of the training, mice had to lick within 3 s after sound offset to receive a drop of soya milk as reward and the trial was considered a correct hit. If animal did not licked within and after the tone trial was considered as missed. If the mice licked while tone was ongoing they received a mild air puff oriented toward the right eye, and a time out (2-3 s) until the next trial could start. These trials were removed from analysis as target (sound offset) could not be correctly delivered. Sounds were delivered without preceding cues at random interstimulus intervals ranging from 3 to 5 s. Licks were detected with a piezo sensor attached to the reward spout. Within consecutive training days, reward window was shortened down to 1 s. *Craniotomy*. Once animals performed at least 30 % of correct hits they were considered initially trained and had a craniotomy performed under ketamine/xylazine (80 mg/kg) and AAF mapping on the same day. *Recordings*. On the following day mice were moved on to a tasting phase where behaviour training was coupled with acute electrophysiology recordings in AAF. During testing phase tones were played at 60 dB SPL and laser was added unilaterally above right-AAF and activated for 200 ms following sound termination in half of the trials. All experiments were performed in a soundproof box (IAC acoustics, Hvidovre, Denmark) and monitored from outside the sound box with a camera (C920, Logitech, Switzerland). Laser power was set around 4.2 mWat and was adjusted every day so it is causing a robust suppression of offset responses in PV-cells. Testing phase was carried out up to 6 days. Behavioral control and data collection were carried out with custom-written programs using a complex auditory processor (RZ6, Tucker Davis Technology, FL, USA), and further analyzed with MATLAB (MathWorks, MA, USA).

## Data Analysis

All data analysis was performed using custom-written MATLAB (2019) (Mathworks) code.

### Tuning receptive fields

To determine BF and tuning receptive fields (TRF) we used PT (50 ms duration, randomized ISI distributed equally between 500 and 1000 ms, 2 repetitions, 4 ms cosine on and 0.01 ms cosine off ramps) varying in frequency from 4 to 48.5 kHz in 0.1 octave increments and in level from 0 to 80 dB SPL in 5 dB increments. Tuning receptive fields, best frequency and spiking rates were calculated in fixed time windows: onset: 6-56 ms, offset: 56–106 ms. TRFs were smoothed with a median filter (4×4 sampling window) and thresholded to 0.2 of peak amplitude. Onset and offset BF was defined as the frequency that elicited maximal response across all sound levels. Onset and offset peak latency was determined as the time point in which the smooth PSTH (kernel=hann (9)) collapsed across all tested stimuli showed maximum response (binning size: 5 ms). Spontaneous activity was calculated based on activity preceding sound onset (150-50 ms, binning size: 5 ms).

### Tone duration responses

To study responses to tones with different durations, we used 10 repetitions of PT (AAF: 9 kHz; MGB: frequency adapted to offset BF of recorded neurons) with 4 ms cosine on and 0.01 ms cosine off ramps, which were varied in duration (50, 100, 150, 250, 350, 500 ms), ISI (gap between 2 stimuli of 50, 100, 250, 500, 1000, 2000 ms) and played at 60 dB SPL. For AAF, frequency was fixed to 9 kHz because 9 kHz PT evoked significant offset responses in almost all tested AAF. Offset spike rates were calculated in a fixed time window of 6-56 ms following sound termination.

### Spectral complexity

To study offset responses in MGB and AAF evoked by sounds with different spectral complexity we recorded responses in both regions to 500 ms pure tones (PT) and white noise (WN) bursts played at 60 dB SPL with 4 ms cosine on and 0.01 ms cosine off ramps (for MGB the PT frequency was chosen based on offset BF of neurons in each session; for the AAF, a fixed PT of 9 kHz was used). WN bursts were not fixed but consisted of randomly chosen noise samples. Offset spike rates were calculated in a fixed time window of 6-56 ms following sound termination.

### Sound rise-fall time study

To study dependence of onset and offset responses on the temporal profile of a tone, we varied rise and fall time at sound onset and offset (0.01, 1, 2, 4, 10, 50, 100, 200 ms). PT (AAF: 9 kHz; MGB frequency adapted to offset BF of recorded neurons) were played at 60 dB SPL for 500 ms and repeated 50 times. Peak amplitude of offset responses was defined in first 100 ms after stimulus onset or offset.

### Offset detection in ongoing sound

To check if offset responses encode changes within ongoing sound we used tone consisting of 3 frequency components (20 kHz, 14 kHz and 9 kHz) played at 60 dB SPL and repeated 60 times. All frequency components had common onset but were terminated at different time points (300 ms, 400 ms, and 500 ms). Offset spike rates were calculated after every component removal in fixed windows: 306-356 ms; 406 – 456 ms; 506 – 556 ms.

### Sound termination detection task

To compare hit rate for trials with and without laser or for different tested ramps a moving average was calculated with window size of 10 trials. The data for each condition was calculated separately. For the average, 10 adjacent trials were taken, and only the behavior corresponding to specific condition were used (meaning that each average is made of a maximum of 10 trials but usually of less). This allows a comparison of performance over time across the tested conditions.

### Decoding population activity

The logistic regression model was used to decode animal performance from neural responses (code from Neuromatch Academy W1D4, www.neuromatchacademy.org). Spontaneous activity (50 – 100 ms before sound onset), onset response (0-50 ms from sound onset), sustained response (500 – 450 ms before sound offset), offset response (0-50 ms from sound offset) or late response (500 – 550 ma after sound offset) were used to train and test the model. L2 regularization was used to avoid over-fitting. 8-fold cross validation was performed by leaving out a random 12,5% subset of trials to test the classifier performance, and remaining trials were used to train the classifier. A range of regularization values was tested (0.0001 to 10000 logspaced), and the one that gave the smallest error on the validation dataset was chosen as the optimal regularization parameter. Classifier accuracy was computed as the percentage of testing trials in which the animal’s choice was accurately predicted by the classifier and summarized as the average across the 10 repetitions of trial subsampling. Spiking activity of each neuron was z-scored before running the logistic regression model. Trial labels were shuffled to confirm that decoding is not working for random data. This procedure was repeated 10 times. Then the average across the 10 repetitions was used to assess classifier accuracy for randomized data. To remove all the sessions with a too small number of trials or too few offset cells, only the sessions with a significant difference classification accuracy between real and shuffled data based on the late response (0.5 s after sound offset) were used.

### Statistical analysis

Statistical tests were performed with GraphPad Prism software version 7.03 (GraphPad Software, USA). The standard error of the mean was calculated to quantify the amount of variation between responses from different populations. Nonparametric, unpaired Mann-Whitney test was used to calculate whether there were any significant differences between medians of recordings in AAF and MGB. Wilcoxon paired test was used to compare differences between paired values obtained in different treatments. Two-way ANOVA was used to test the main effects of sound duration and intervals on offset responses and their interaction effect. Dunn’s multiple comparisons test was used to perform pairwise multiple comparisons. Spearman correlation tests were used to test for significant associations between pairs of variables measured with ranking.

## ACKNOWLEDGEMENTS

This work was supported by grants from the Lundbeck Foundation (Fellowship to T.R.B) and the Swiss National Science Foundation (ERC Transfer grant to T.R.B). We thank Gioia De Franceschi for help with data analysis and Boris Gourévitch for thoughtful comments on the manuscript.

## AUTHOR CONTRIBUTIONS

M.S. and T.R.B. designed the study and wrote the manuscript; M.S performed all experiments and analysed the data.

