## Supplementary figures for "Emergence and function of cortical offset responses in sound termination detection"

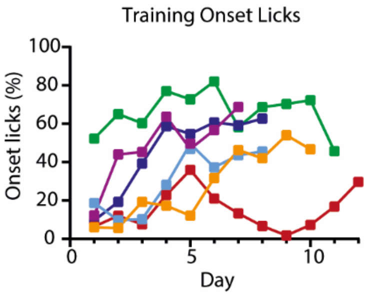


**Figure S1. Occurrence of offset licks during training. Related to Figure 1.**

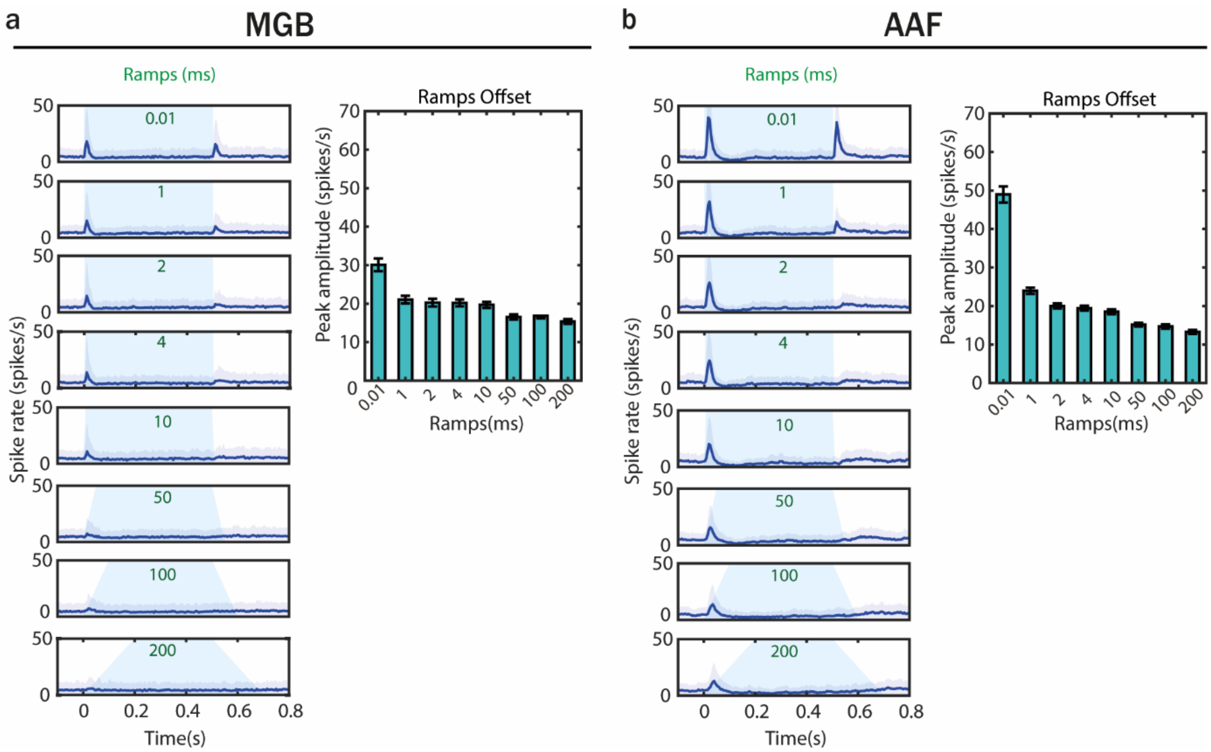


**Figure S2. Offset responses evoked by sounds terminated with different fall ramps emerge already in MGB. Related to Figure 2.**

(**a**) (Left) averaged PSTH (mean± STD) of MGB neuron's response to PTs (9 kHz) played at 60 dB SPL with varied on and off ramps: 0.01, 1, 2, 4, 10, 50, 100, 200 ms. (Right) comparison of offset peak amplitude evoked by PTs with different onset and offset ramps. (**b**) (Left) averaged PSTH of AAF neuron's response to PTs (frequency adapted to offset BF of recorded neurons) played at 60 dB SPL with varied on and off ramps. (Right) comparison of offset peak amplitude evoked by PTs with different onset and offset ramps.

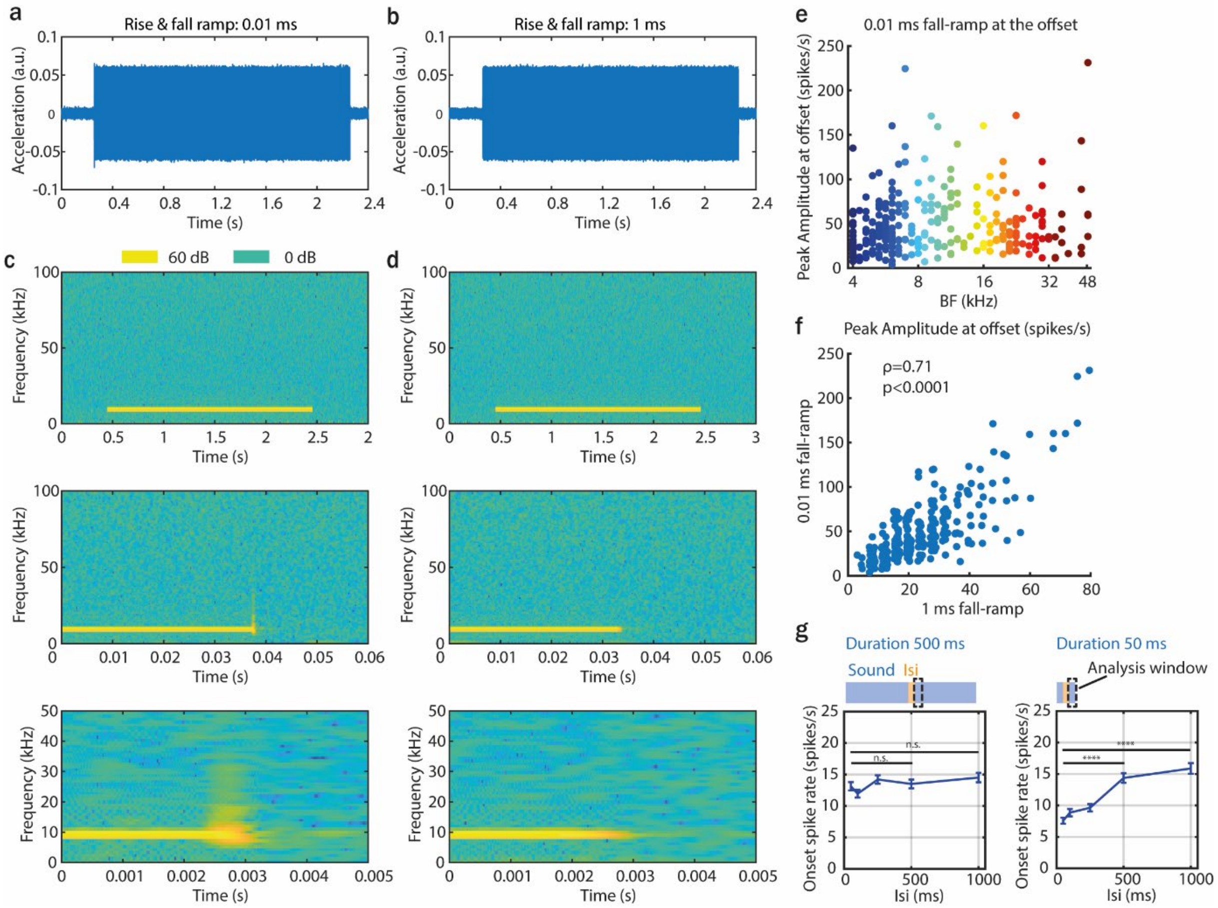


**Figure S3. Properties of 9 kHz pure tones played at 60 dB SPL with different rise and fall-ramps. Related to Figure 2.**

(**a, b**) Acceleration traces of 9 kHz pure tone played at 60 dB SPL with 0.01 ms (a) and 1 ms (b) rise and fall-ramp measured with an ultrasensitive microphone (Avisoft-Bioacoustics CM16/CMPA). (**c, d**) Spectrograms of a 9 kHz pure tone played at 60 dB SPL with 0.01 ms (c) and 1 ms (d) rise and fall-ramp. (**e**) Peak amplitude of offset responses evoked by 0.01 ms fall ramp as a function of onset best frequency. (**f**) Comparison of peak amplitude of offset responses evoked by 0.01 ms and 1 ms fall ramp (ρ=0.71, p<0.0001, Spearman correlation). (**g**) Modulation of onset responses by preceding offset (left) or onset (right) responses.

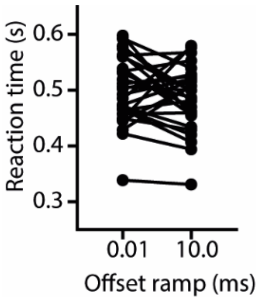


**Figure S4. Comparison of reaction times in sound termination detection task for two tested fall-ramps: 0.01 ms and 10.0 ms. Related to Figure 2.**

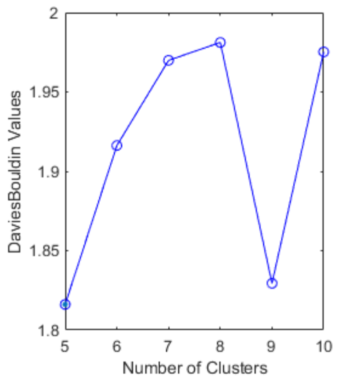


**Figure S5. Davies-Bouldin index for different number of clusters. The minimum value is reached for 5 and 9 clusters. Related to Figure 4.**

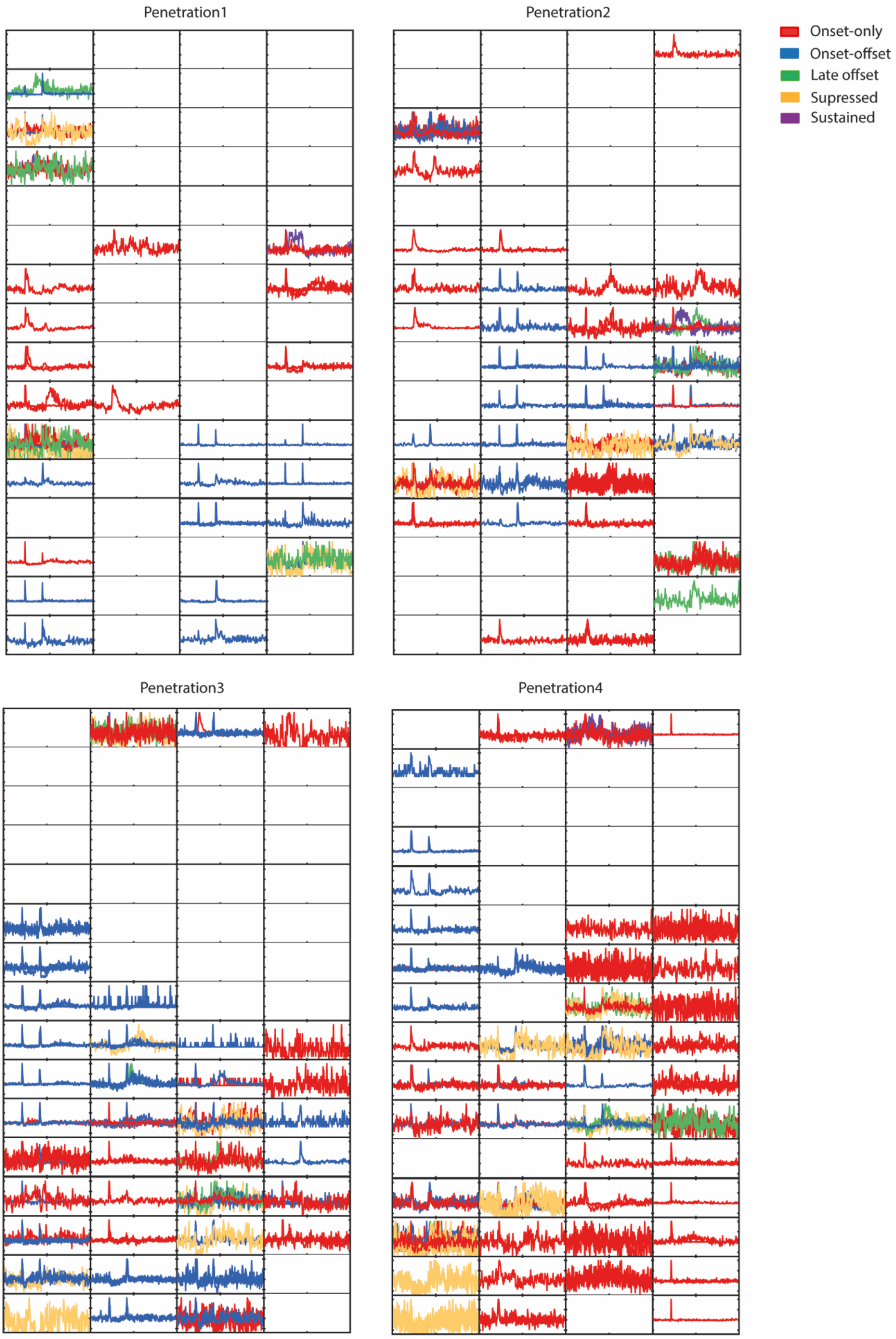


**Figure S6. 2D representation of temporal dynamic of MGB cells recorded during 4 experiments with 64-channel electrode (16x4). Related to Figure 4.**

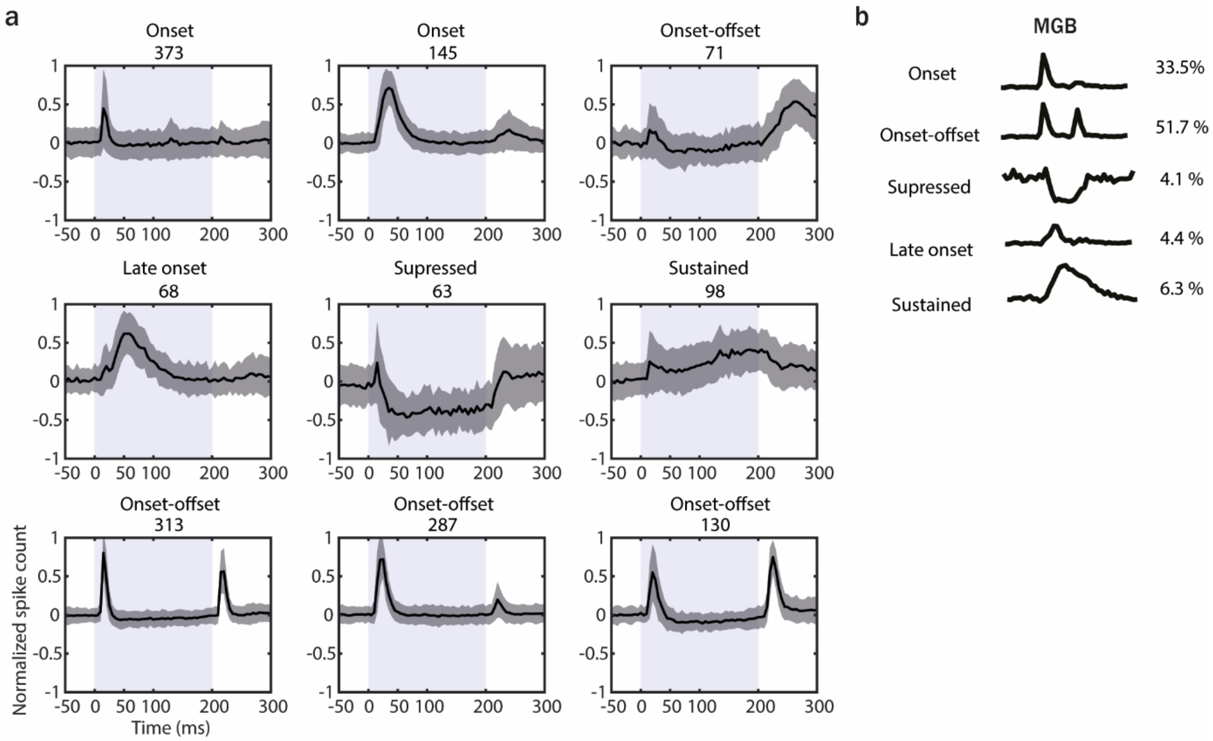


**Figure S7 Response dynamics of MGB cells recorded during antidromic stimulation experiment.ts. Related to Figure 5**.

(**a**) Results of k-means clustering performed on MGB cells recorded during antidromic stimulation experiments (n=1548) evoked by 200 ms PT with varied frequency (4 to 48.5 kHz) and sound level (0 to 80 dB SPL) presented with randomized ISI (500–1000 ms). The graphs represent the mean temporal dynamic of cells belonging to each cluster. Data represent mean ± SD. The grey shaded bars represent the tone. (**b**) Percentage of cells with distinct temporal dynamic of responses in the MGB cells recorded during antidromic experiment.

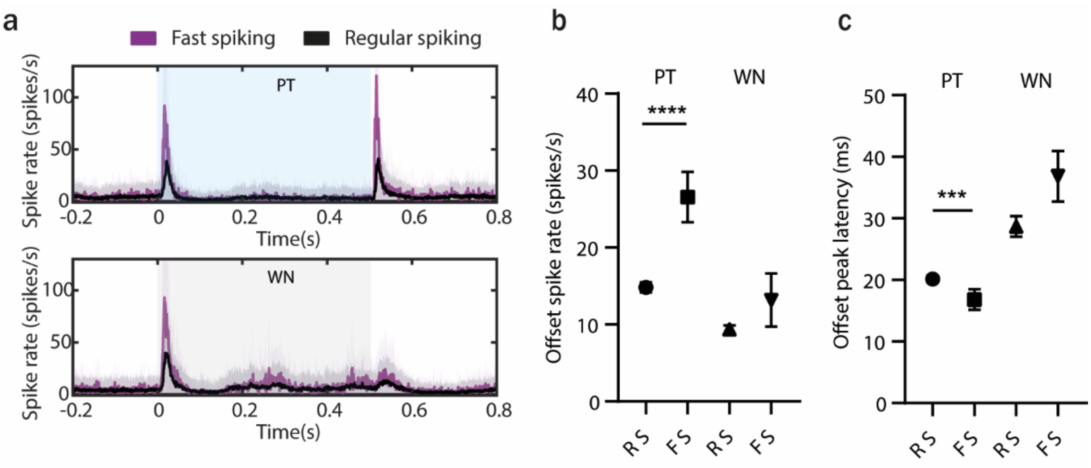


**Figure S8. Offset processing in fast and regular spiking AAF neurons. Related to Figure 7.**

(**a**) Averaged PSTH (mean±STD) of fast and regular spiking AAF neuron's response to PTs (9 kHz) or WN bursts played at 60 dB SPL with sound duration of 500 ms and ISI between 500 and 2000 ms. (**b**) Comparison of offset spike rate for RS and FS units in response to termination of 500 ms PT or WN, p<0.0001, Mann-Whitney test. (**c**) Comparison of offset response latencies of RS and FS units in response to termination of 500 ms PT or WN, p=0.0006, Mann-Whitney test.

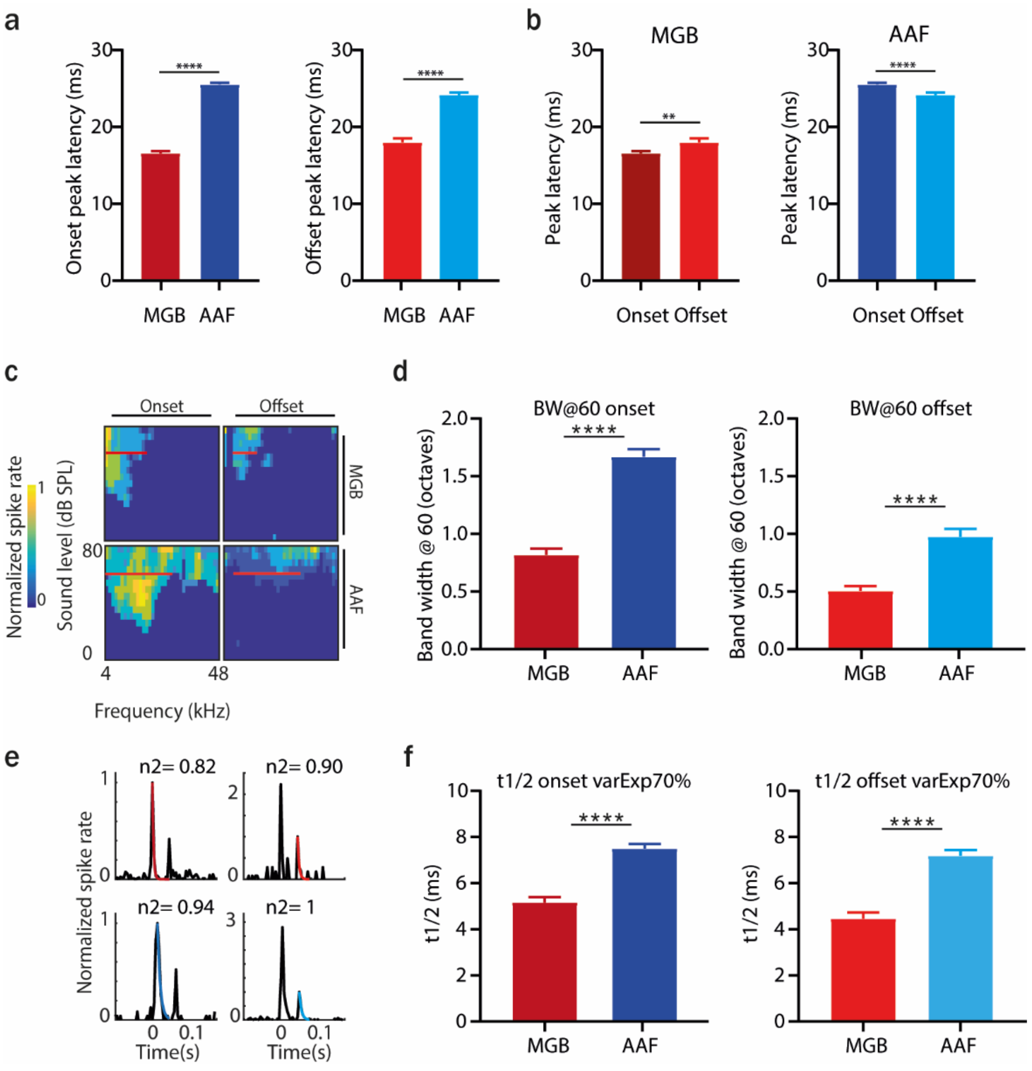


**Figure S9**. **Distinct spectral and temporal tuning properties of MGB and AAF cells.** (**a**) Comparison of onset (left) and offset (right) peak latency between MGB and AAF neurons. Data represent mean ± SEM. (**b**) Comparison of onset and offset peak latency within MGB (left) and AAF (right) neurons. Data represent mean ± SEM. (**c**) Example of tuning receptive fields (TRF) of onset and offset responses of an MGB and AAF neuron. (**d**) Comparison of tuning band width at 60 dB SPL of onset and offset responses in MGB and AAF neurons. (**e**) Example fit of exponential decay model to PSTH of MGB and AAF neurons' onset and offset responses (n2: amount of variance explained). (**f**) Comparison of half decay time (t1/2) of onset and offset responses in MGB and AAF neurons. Data represent mean ± SEM.
